# The uplift of the Qinghai-Tibetan Plateau reshaped genome architecture and diversification of Asian torrent catfishes

**DOI:** 10.64898/2026.09.30.755532

**Authors:** Liandong Yang, Yuting Qian, Minghui Meng, Tingru Mao, Wenjun Chen, Zufa Ding, Yang Liu, Lingling Zheng, Cheng Wang, Xiaoni Gan, Yiyu Chen, Shunping He

## Abstract

Tectonic uplift and mountain building are major drivers of species diversification and adaptation, yet how they reshape genome architecture and promote evolutionary innovation remains poorly understood. Here, we generated multiple chromosome-level genomes for Asian torrent catfishes (Sisoridae) and performed phylogenomic and comparative genomic analyses. Our results support an Asian origin of Sisoridae and provide a refined phylogenetic topology for catfishes. Comparative analyses revealed extensive chromosomal rearrangements during sisorid evolution, indicating highly dynamic genome architectures. The timing of sisorid diversification overlaps with major phases of regional tectonic change, and analyses of phylogenetic discordance indicate heterogeneous contributions from incomplete lineage sorting and introgression. Elevational distribution was associated with variation in estimated molecular evolutionary rates, while exploratory scans identified lineage-specific candidates for selection and convergent evolution. Candidate coding and non-coding changes occur near genes with known roles in fin development, body patterning, and liver biology, but their effects on sisorid traits remain to be tested. Together, our results demonstrate how tectonic uplift reshaped genome architecture and diversification in Asian torrent catfishes and provide new insights into the genomic basis of adaptation in mountainous freshwater ecosystems.

## Introduction

The family Sisoridae is one of the most diverse catfish lineages in Asia, and its taxonomy and evolutionary history have attracted considerable attention(1, 2). Early classifications based primarily on external morphology and osteological characteristics divided Sisoridae into the subfamilies Sisorinae and Glyptosternae, with glyptosternoid fishes generally corresponding to the latter group (3, 4). Both Sisoridae and glyptosternoids have traditionally been regarded as monophyletic based on shared morphological synapomorphies, particularly traits associated with adaptation to fast-flowing habitats(5–7). However, molecular phylogenetic analyses employing mitochondrial genomes and limited nuclear markers have yielded discordant topologies, challenging the monophyly of these groups(8, 9). Resolving these phylogenetic uncertainties is essential for determining whether the remarkable morphological diversity of Sisoridae represents shared evolutionary history or repeated adaptive evolution.

Members of the family Sisoridae exhibit a remarkable repertoire of morphological modifications associated with torrent adaptation, including dorsoventrally flattened bodies, enlarged paired fins, accessory livers, and specialized adhesive apparatuses that enable attachment to rocky substrates in rapid currents(10–12). However, whether this diversity represents a single adaptive radiation or repeated convergent evolution driven by similar hydrodynamic pressures remains unresolved(13, 14). Glyptosternoid fishes represent one of the most specialized lineages within Sisoridae; in addition to torrent-adapted traits, they are also widely regarded as a lineage adapted to high-altitude environments. Distributed primarily across the Qinghai-Tibetan Plateau and adjacent mountainous regions, these fishes experience extreme conditions characterized by low temperatures, hypoxia, and high ultraviolet radiation(15, 16). Therefore, Sisoridae provides an excellent system for investigating how ecological constraints and geological processes interact to shape adaptive evolution in extreme environments.

Siluriformes is among the most species-rich and geographically widespread groups of Teleostei fishes, exhibiting extensive ecological and morphological diversification across global freshwater ecosystems. However, deep relationships among major siluriform lineages remain debated due to conflicting signals from morphological, mitochondrial, and phylogenomic datasets(17–20). Uncertainty regarding interfamilial relationships and the monophyly of several groups (Amphiliidae and Mochokidae) has hindered reconstruction of catfish diversification history and obscured the evolutionary origin of specialized lineages such as Sisoridae(21–23). The diversification of Sisoridae is thought to be closely associated with the uplift of the Qinghai–Tibetan Plateau, which profoundly reshaped Asian topography, climate, and river systems during the Cenozoic(24, 25). These environmental changes may have promoted lineage diversification through geographic isolation and ecological opportunity(26–28). However, testing the evolutionary relationship between plateau uplift and Sisoridae diversification requires a well-resolved phylogeny of Siluriformes, combined with temporal and biogeographic reconstruction.

Adaptive radiation in extreme environments is frequently accompanied by ecological specialization and complex evolutionary processes. The Qinghai–Tibetan Plateau represents a major biodiversity hotspot where endemic lineages have evolved diverse adaptations to hypoxia, temperature extremes, and other environmental challenges(29–31). Increasing evidence indicates that incomplete lineage sorting (ILS), introgression, and hybridization can substantially influence diversification trajectories(32–35). Previous studies of plateau organisms have revealed molecular mechanisms underlying adaptation, including changes associated with oxygen metabolism, energy regulation, and physiological remodeling(36–41). Nevertheless, the genomic basis of adaptation in Sisoridae catfishes remains poorly characterized, particularly regarding the contributions of genome structural evolution, gene family diversification, convergent evolution, and lineage-specific molecular adaptation.

Recent advances in long-read sequencing technologies and comparative genomics have dramatically enhanced our ability to resolve complex evolutionary histories and uncover the genomic basis of diversification and environmental adaptation. Chromosome-level genome assemblies enable detailed investigations of genome evolution, phylogenetic conflict, introgression, and adaptive genomic changes. In this study, we generated chromosome-level genome assemblies for multiple representative Sisoridae catfishes and assembled a comprehensive phylogenomic dataset spanning Siluriformes. Using comparative genomic and phylogenomic approaches, we reconstructed the evolutionary origin and phylogenetic relationships of Sisoridae and glyptosternoid catfishes within a broad siluriform context. We further investigated diversification dynamics, phylogenetic conflict, incomplete lineage sorting, and introgression associated with Sisoridae radiation. Additionally, we explored the genomic basis of adaptation through analyses of chromosome evolution, gene family evolution, positive selection, and convergent evolution. Together, our study provides a comprehensive genomic foundation for understanding the origin, diversification, and adaptive evolution of East Asian endemic Sisoridae catfishes.

## Result

### Origin of Sisoridae and phylogenetic reconstruction of Siluriformes

Sisoridae represents a distinctive lineage of benthic catfishes characterized by remarkable adaptations to fast-flowing riverine environments in East Asia. However, its phylogenetic placement and biogeographic origin remain contentious, as morphological resemblance to African amphiliid catfishes has led to competing hypotheses of African versus Asian origins, while its subsequent radiation in Asia has been proposed to coincide with the uplift of the Qinghai-Tibet Plateau(21). To resolve the evolutionary history of Sisoridae in the context of Siluriformes evolution, we reconstructed the relationships of Siluriformes using genome and transcriptome data from 181 species: 180 Siluriformes representing 28 major families plus one outgroup species. This dataset includes newly generated genomes from nine Sisoridae species, two Amblycipitidae species, three Amphiliidae species and one Mochokidae species (Supplementary Fig 1).

We reconstructed the phylogeny of Siluriformes using 1,896 BUSCO orthologues based on multiple datasets and inference strategies, including concatenated matrices of all codon positions, first and second codon positions, third codon positions, rate-stratified gene sets (fast-evolving group 1 and slow-evolving group 2), and a coalescent-based species tree inferred using ASTRAL from individual gene trees. (Supplementary Fig 2-8). At the family level, the topology inferred from the all-codon dataset was highly congruent with that derived from third-codon positions and the fast-evolving gene set (group1). These three datasets recovered nearly identical relationships among major siluriform lineages, with only minor topological discrepancies observed among Amphiliidae, Mochokidae, and Malapteruridae. The ASTRAL species tree was also broadly consistent with the all-codon topology in the overall phylogeny of Siluriformes. However, differences were observed in the placement of Plotosidae, Chacidae, Clariidae, and Siluridae. Notably, these conflicting nodes were associated with low local posterior probability (LPP) values in the ASTRAL analysis, suggesting weak phylogenetic support. In contrast, the all-codon tree and rate-based concatenated analyses consistently recovered these relationships with higher support. Based on the overall topological congruence, we selected the all-codon topology as the final species tree (Figure 1A). Across the phylogeny, multiple short internal branches were observed, particularly at deeper nodes within Siluriformes, suggesting a history of rapid diversification(42–44).

**Figure 1.**
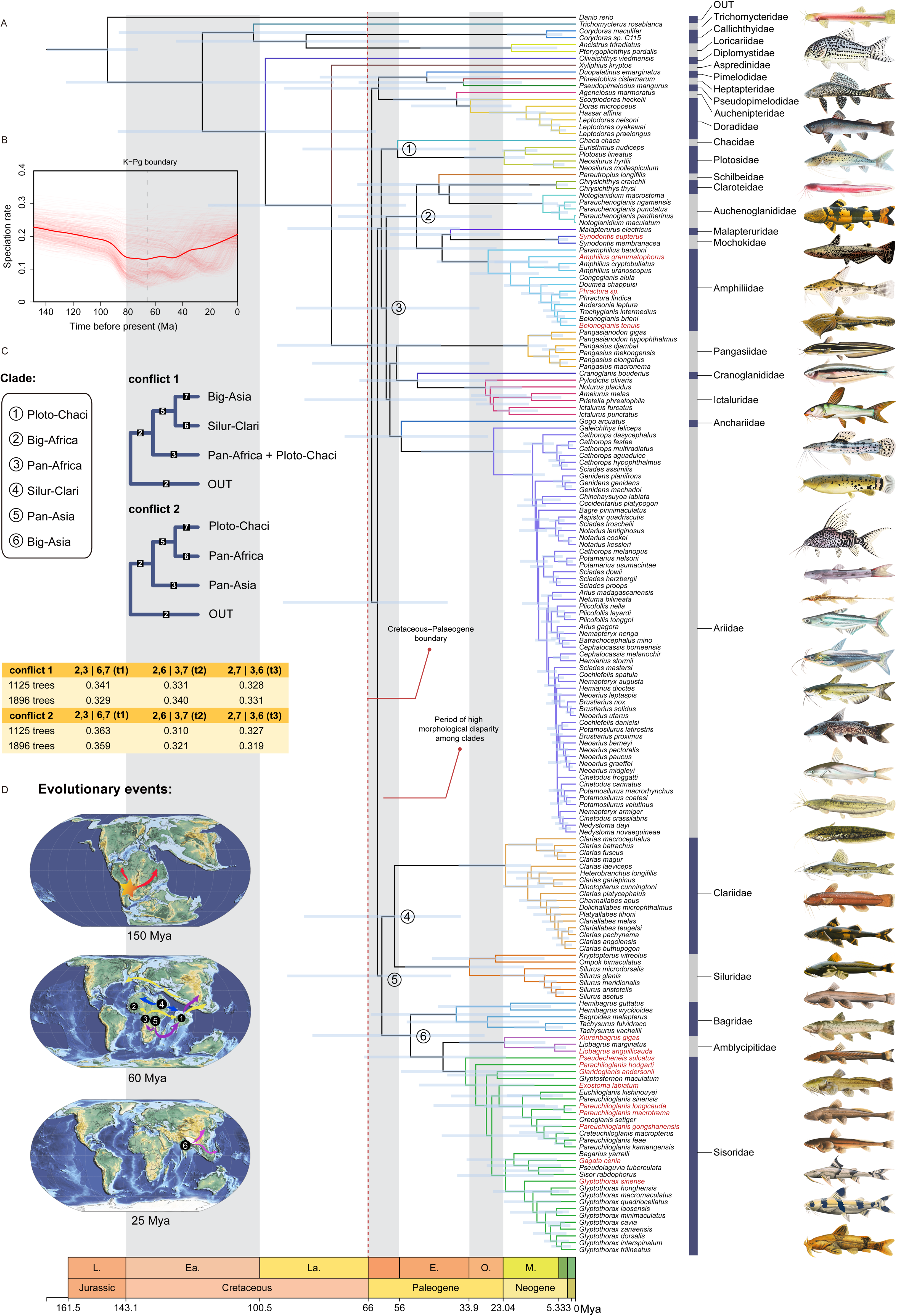
phylogenetic reconstruction of Siluriformes. Maximum-likelihood (ML) phylogeny of Siluriformes reconstructed using BUSCO genes with 90% taxon occupancy, including 181 species in total: 180 Siluriformes representing 28 families and one outgroup species. Most major nodes received high bootstrap support values. Light blue rectangles at internal nodes indicate the 95% confidence intervals of estimated divergence times. Numbers 1-6 in circles denote the six major clades of Siluriformes. Species names shown in red represent genomes newly generated in this study. The red dashed line indicates the Cretaceous-Paleogene extinction event boundary. **B.** Rate-through-time plots show posterior diversification dynamics of sisorid catfishes, where thin red lines represent up to 500 randomly sampled posterior MCMC samples (smoothed), and the thick red line indicates the mean speciation rate through time (smoothed). **C.** Names of the six major clades in the phylogenetic tree of Siluriformes, and the two major types of phylogenetic conflict along with the proportions of their alternative topologies. **D.** Evolutionary events in Siluriformes: Late Jurassic (∼150 Ma), breakup of western Gondwana with South America still connected to Africa, corresponding to the inferred South American origin and initial diversification of Siluriformes. Early Cenozoic (∼60 Ma), post-K-Pg continental reorganization and northward drift of the Indian plate, corresponding to a major diversification node within Siluriformes with pronounced phylogenetic discordance and ILS; yellow and purple arrows indicate Pan-Africa and Pan-Asia lineages, and blue arrows indicate the Silur-Clari divergence. Neogene (∼25 Ma), uplift of the Qinghai–Tibet Plateau, corresponding to diversification of the Asian “big-Asia” clade.

Following previous studies, we retained the “Big Asia” and “Big Africa” biogeographic groupings within Siluroidei(17, 19). The topology of the Big Asia clade is highly stable across all analyses, consistently comprising Bagridae, Amblycipitidae and Sisoridae. Notably, Sisoridae is always recovered as the sister group to Amblycipitidae, supporting an Asian origin of Sisoridae rather than an African one. In contrast, relationships within the Big Africa clade differ from previous studies, particularly in the placement of Mochokidae and Amphiliidae. By incorporating newly sequenced genomes, we resolve these inconsistencies and recover a monophyletic group comprising Mochokidae, Amphiliidae and Malapteruridae. We further extend this classification by defining two broader groupings: Pan-Asia, which includes the Asian-origin lineage and its sister clade, the Silur-Clari lineage (Siluridae and Clariidae), although several descendant lineages remained and diversified in Africa, and Pan-Africa, which encompasses several families now distributed in Southeast Asia, the Americas and Australia, reflecting a pan-African origin (Supplementary Fig 9). These results provide a new perspective on the evolutionary history of Siluriformes.

Across all phylogenetic reconstructions of Siluroidei, the placements of the Silur-Clari and Ploto-Chaci clades remain the most difficult to resolve. To further investigate these uncertainties, we constructed gene trees using orthologous datasets with 90% and 85% taxon occupancy (1125 and 1896 genes, respectively) and quantified topological heterogeneity among loci. Despite the general consistency of the species-tree results, two major topological conflicts were identified: one concerning whether the Silur-Clari clade forms a monophyletic group with the Big Asia clade, and the other involving the relationship between the Ploto-Chaci clade and the Pan-Africa clade (Figure 1C). These conflicts likely reflect rapid early divergences and incomplete lineage sorting during the radiation of Siluroidei, and may underlie the persistent difficulty in resolving the phylogenetic positions of the Silur-Clari and Ploto-Chaci lineages.

We next estimated divergence times using 12 fossil calibration points (Supplementary Table 2). The results suggest that Siluriformes originated in South America approximately 158 million years ago and established its primary lineage structure during the Cretaceous. The major clades Pan-Asia, Pan-Africa and Ploto-Chaci diverged between ∼66 and 56 million years ago. The rate-through-time analysis revealed a bottleneck and a subsequent marked increase in diversification rate in Siluriformes at the Cretaceous-Palaeogene (K-Pg) boundary (∼66 Ma). (Figure 1B).

This elevated diversification signal persists throughout the early Paleogene, coinciding with the initial radiation of major lineages including the Pan-Asia, Pan-Africa, and Ploto-Chaci clades. Similar post-K–Pg diversification patterns have also been reported in multiple vertebrate lineages(45–47). The Big-Asia lineage within Pan-Asia, along with most representatives of Siluridae and several Clariidae lineages, subsequently colonized Asia through the Indian Plate stepping-stone route and experienced subsequent diversification(48). Most family-level divergences within this clade occurred between ∼40 and 20 million years ago, coinciding with the Cenozoic India– Asia collision and the uplift of the Qinghai–Tibet Plateau(49). These temporal patterns indicate a close association between geological events and lineage diversification in Asian catfishes (Figure 1D).

Together, these results suggest that the origin and diversification of Siluriformes were shaped by tectonic movements and climatic transitions, with intercontinental dispersal facilitated by land bridges and oceanic changes. Within this context, the Asian origin and subsequent radiation of Sisoridae provide a basis for investigating their adaptive diversification in fast-flowing riverine environments.

### Genome sequencing and comparative genomics of Sisoridae

To comprehensively investigate genome evolution and structural diversity in Sisoridae, we generated and analysed a new comparative genomic dataset comprising nine species. In this study, we newly assembled genomes for nine Sisoridae species, among which seven were assembled at the chromosome level (Supplementary Table 3). We performed comprehensive genome annotation, including both protein-coding genes and repetitive elements (Supplementary Table 4). In addition, three previously published Sisoridae genomes were incorporated into downstream comparative genomic analyses.

Sisoridae exhibits remarkable genomic and karyotypic diversity, with genome sizes ranging from 478 (*Gag. cenia*) to 900 (*Pare. macrotrema*) Mb and chromosome numbers varying from 2n=44 (*Par. hodgarti*) to 2n=60 (*Glar. andersonii*) (Figure 2A and Supplementary Table 3). In addition, repeat content differs substantially among lineages, with species of the genus *Pareuchiloglanis* showing particularly high proportions of repetitive elements.

**Figure 2.**
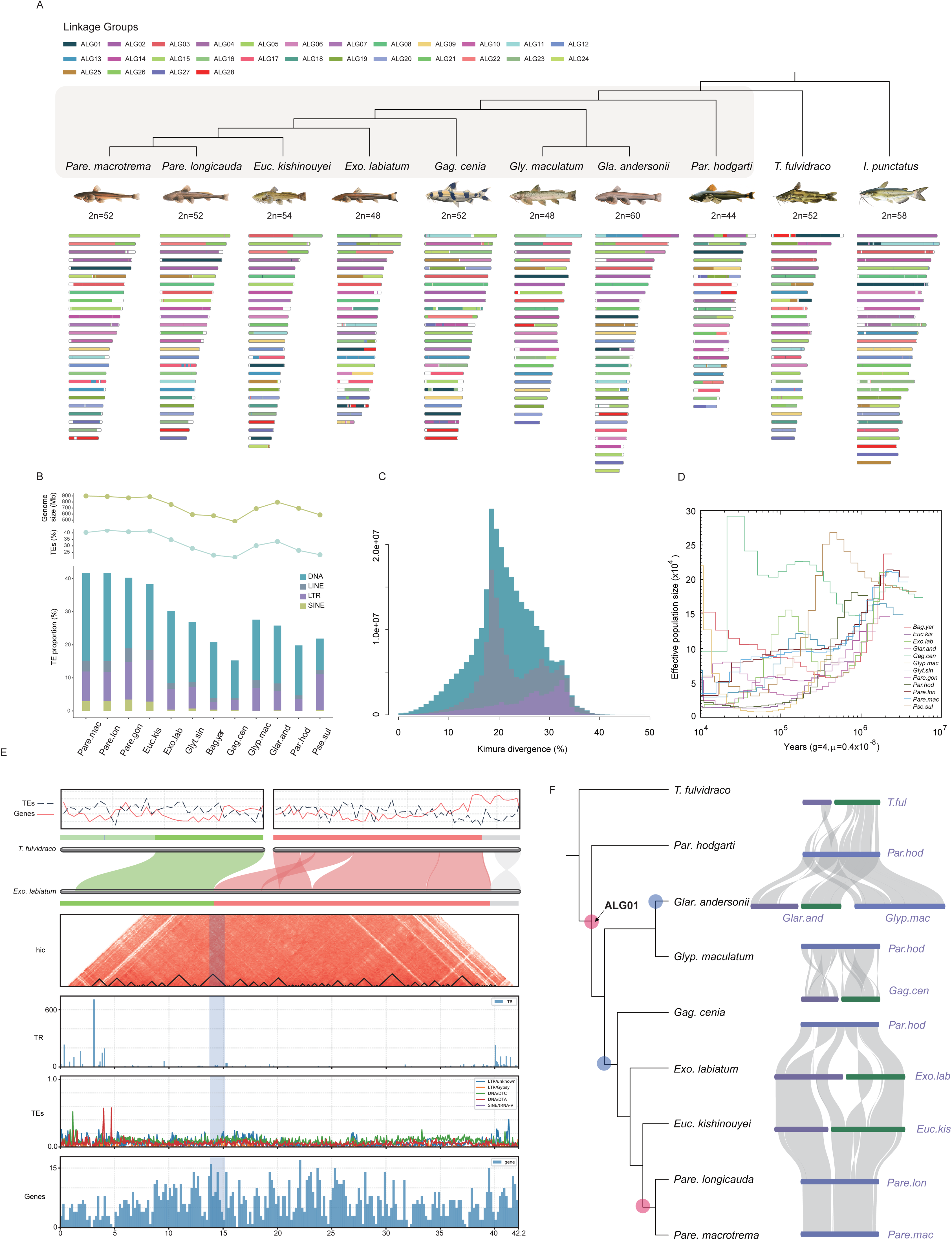
Ancestral linkage groups (ALGs) and chromosomal evolution of Sisoridae catfish. **A.** The fraction of genes derived from each ALGs (depicted as squares named ALG01-28) in various catfish species, including 8 Sisoridae catfish. **B.** Genome size, TE size, and TE type content in Sisoridae catfish. **C.** TE landscape plots obtained using Kimura distance-based copy divergence analyses showing the expansion patterns of TEs in Sisoridae species, with *Exo. labiatum* shown as a representative; results for the remaining species are provided in Supplementary Fig 10. **D.** Changes in effective population size (Ne) over the past three million years inferred using the PSMC method. **E.** The syntenic relationships between chr3 of *Exo. labiatum* (formed by the fusion of ALG22 and ALG16) and the corresponding chromosomes in *T. fulvidraco*. From top to bottom, the genomic tracks represent: gene distribution (black line) and TE density (red line) along the chromosomes of *T. fulvidraco*; syntenic alignments between chr2 and chr10 of *T. fulvidraco* and chr3 of *Exo. labiatum*; Hi-C interaction signals and predicted TAD structures on chr3 of *Exo. labiatum*; tandem repeat (TR) distribution; TE distribution; and protein-coding gene distribution. ALG01 underwent multiple rounds of chromosomal fission and fusion across different branches of the Sisoridae phylogeny. Pink circles at the nodes indicate fusion events, whereas blue circles indicate fission events.

To investigate the genomic basis underlying this diversity, we generated and analysed high-quality genome assemblies from six newly sequenced Sisoridae species, together with two previously published chromosome-level genome. Based on these datasets, we reconstructed ancestral linkage groups (ALGs) for Sisoridae (Figure 2A). Genome-wide synteny analyses based on the reconstructed ALGs further demonstrated that, despite extensive conservation of ancestral chromosomal identity across Sisoridae genomes, numerous lineage-specific disruptions of synteny were evident. Comparisons with the reconstructed ancestral genome identified multiple Dx and Dy peaks corresponding to putative chromosomal rearrangement breakpoints, providing independent support for pervasive structural reorganization during Sisoridae chromosome evolution (Supplementary Fig 11 and 12). Our analyses revealed extensive chromosomal rearrangements in Sisoridae compared with the relatively conserved haploid chromosome number (n=26-29) observed across other Siluriformes lineages(50–53). Notably, *Par. hodgarti*, a basal lineage within Sisoridae, shows extensive chromosomal fusions relative to its closest non-sisorid sister lineage (*Tachysurus fulvidraco*), indicating early and substantial karyotypic restructuring during Sisoridae evolution. In contrast, members of the genus *Pareuchiloglanis* largely retain a relatively conserved karyotype (2n=52) and exhibit high levels of genome-wide synteny, highlighting lineage-specific heterogeneity in chromosomal evolutionary dynamics within Sisoridae.

We next examined genome size variation and transposable element (TE) content across Sisoridae species. Genome size was strongly positively correlated with TE abundance, indicating that TE expansion represents a major driver of genome size evolution in this lineage (Figure 2B). Among TE classes, DNA transposons contributed most significantly to genome expansion, as also observed in most teleost fishes (Supplementary Table 5)(54). To further investigate TE evolutionary dynamics, we performed Kimura divergence-based analyses of TE landscapes (Figure 2C). Across Sisoridae species, we observed broadly consistent patterns of TE accumulation. TE expansion occurred in a stepwise manner: LTR retrotransposons exhibited the earliest expansion, followed by two distinct waves of LINE activity, whereas DNA transposons expanded most recently. This indicates a temporally ordered activation of different TE classes during Sisoridae evolution.

We further inferred historical effective population size (Ne) changes over the past ∼3 million years using PSMC analyses (Figure 2D). We observed a notable association between genome size and demographic history: species with smaller effective population sizes (e.g. *Pare. macrotrema* and *Pare. longicauda*) tend to possess larger genomes, whereas species with larger Ne (e.g. *Gag. cenia*) generally maintain smaller genomes. This pattern indicates a potential link between genome size variation and demographic history in Sisoridae.

We next investigated chromosomal structural variation in Sisoridae. Several ancestral chromosomal fusion events are widely conserved across species, including the fusion of ALG22 and ALG16, which is shared by most Sisoridae lineages but absent from *Gag. cenia* and *Pare. hodgarti* (Supplementary Fig 13). For example, in *Exo. labiatum*, chromosome 3 (derived from this fusion) shows clear syntenic correspondence with *T. fulvidraco* chromosomes (Figure 2E). Genome-wide tracks integrating gene density, TE distribution, tandem repeats (TRs), Hi-C interaction signals, and predicted TAD structures further reveal that TR enrichment and local chromatin interaction patterns mark remnants of ancestral centromeric regions at fusion breakpoints. Meanwhile, novel TAD structures appear to have formed at these regions, indicating that chromosomal fusion has reshaped three-dimensional genome organization.

We further observed repeated cycles of chromosomal fission and fusion in ALG01 across multiple Sisoridae lineages (Figure 2F). Specifically, relative to outgroup species, ALG01 originated from the fusion of two ancestral chromosomes, followed by independent fission events in the *Par. hodgart*i lineage and in the ancestral lineage of *Gagata*, *Exostoma*, and *Pareuchiloglanis*, and was subsequently fused again at the ancestral node of the genus *Pareuchiloglanis*, highlighting highly dynamic chromosomal remodeling during diversification.

### Genomic diversity during Sisoridae diversification is shaped by incomplete lineage sorting and introgression

To obtain a robust and well-resolved phylogeny of Sisoridae, we conducted a genome-wide phylogenomic analysis. Gene trees were first inferred from filtered WGA blocks using IQ-TREE, and a coalescent-based species tree was subsequently reconstructed in ASTRAL, complemented by a genome-wide assessment of gene tree discordance. The WGA-based phylogeny rejects the monophyly of Glyptosternoid fishes. Taxa traditionally classified as non-glyptosternoids (e.g., *Gagata*, *Bagarius*, and *Glyptothorax*) form a clade that is sister to the lineage comprising representative glyptosternoid genera (e.g., *Euchiloglani*, *Exostoma*, and *Pareuchiloglanis*). Within Sisoridae, *Pseudecheneis* is consistently recovered as the earliest diverging lineage, whereas *Par. hodgarti*, a monotypic genus with multiple derived adaptive traits, occupies a distinct phylogenetic position, suggesting a lineage-specific evolutionary history (Figure 3A). These patterns indicate that similar but not identical adaptive features observed across sisorid fishes likely evolved independently in different lineages, with some traits potentially arising through convergent evolution.

**Figure 3.**
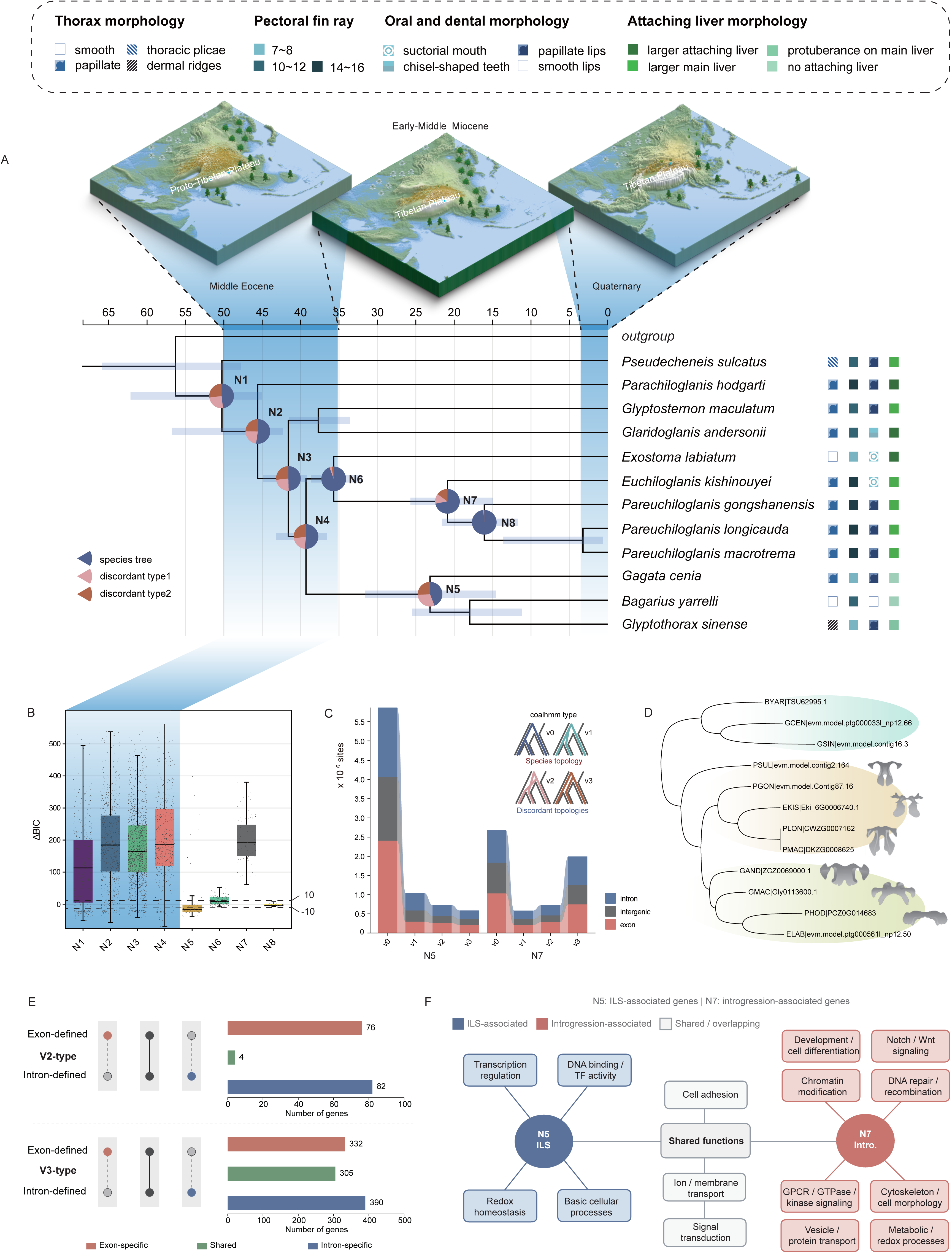
Phylogenetic reconstruction of Sisoridae reveals introgression and incomplete lineage sorting (ILS) as joint sources of genetic diversity during diversification. **A.** Phylogenetic tree of Sisoridae reconstructed based on whole-genome alignment (WGA) data, revealing extensive topological discordance within the family. Pie charts at nodes indicate the support proportions for alternative topologies. Divergence times are shown in the context of major phases of Tibetan Plateau uplift. Paleogeographic reconstructions associated with different uplift stages were adapted from Lu et al. (2023). Colored blocks with distinct patterns denote morphological traits associated with adaptation to plateau torrential habitats across Sisoridae species. **B.** Distribution of ΔBIC values across different nodes in the Sisoridae phylogeny inferred using QuIBL analysis. Each point represents an individual test group, while the boxplots summarize the overall distribution trends of ΔBIC values across all test groups. Black dashed lines indicate the threshold values of −10 and 10. **C.** COALHMM-based estimates of N5 and N7 showing the distribution of v0–v3 sites (species-tree and alternative topological states) and their proportions across genomic regions, presented as stacked plots. **D.** Clustering patterns of *hpxa* show incongruence with species phylogeny, instead grouping more consistently with accessory liver morphology. **E.** Comparison of exon- and intron-defined V2- and V3-type introgression-associated genes. V2- and V3-type genes were identified independently using exon or intron sites. Numbers indicate genes uniquely identified from exon or intron regions, as well as genes shared between the two datasets. V2-type genes exhibited minimal overlap between exon- and intron-derived gene sets, whereas V3-type genes showed substantially greater concordance across coding and non-coding regions. **F.** Functional annotation diagram of genes associated with incomplete lineage sorting (ILS) and introgression.

To disentangle the evolutionary processes underlying phylogenetic discordance, we evaluated the relative contributions of incomplete lineage sorting (ILS) and introgression based on node-specific signals inferred from QuIBL and genome-wide patterns recovered by AutoCoalHMM. Early-diverging nodes (N1-N4) showed a stronger contribution from introgression, with a secondary signal of ILS. This signal coincides with the early uplift stages of the Qinghai-Tibetan Plateau, during which rapid paleogeographic restructuring likely promoted intermittent connectivity among lineages(25, 55). According to the node-level summary of QuIBL model support (Supplementary Table 7), later-diverging nodes exhibited contrasting patterns, with node N5 classified as ILS-dominated, node N7 as introgression-dominated, and nodes N6 and N8 remaining inconclusive because fewer than half of the discordant triplets provided significant model support (|ΔBIC| > 10) (Figure 3B). Consistent with these classifications, the estimated introgression proportion (totalIntroProp) was substantially higher for the introgression-dominated nodes than for the ILS-dominated node, with early-diverging nodes (N1-N4) generally exhibiting the strongest introgression signals across QuIBL triplets (Supplementary Fig 14).

Among the later-diverging nodes, node N5 and nodeN7 were selected as representative examples of ILS- and introgression-dominated discordance, respectively, because they exhibited strong and consistent support for contrasting evolutionary processes across significant QuIBL tests (Supplementary Table 7). We further examined the genomic distribution of these signals using CoalHMM-derived state assignments. Introgression-associated signals were broadly distributed across the genome and more frequently retained across large genomic regions, whereas ILS-associated signals were more fragmented and showed a more transient genomic distribution (Figure 3C and Supplementary Table 8). These differences suggest distinct genomic persistence of the two processes, with introgressed variation contributing more continuously to present-day genomic diversity. To explore the functional characteristics of genes associated with different sources of genealogical discordance, we compared the functional annotations of genes identified from the ILS-dominated (node N5) and introgression-dominated (node N7) analyses (Supplementary Table 9-10). Although no Gene Ontology category remained significantly enriched after multiple-testing correction, clear differences in functional composition were observed between the two gene sets (Figure 3F). Genes associated with node N5 were distributed across diverse fundamental cellular functions, including transcriptional regulation, ion transport, membrane transport, cell adhesion, and redox homeostasis. In contrast, genes associated with node N7 were predominantly involved in developmental regulation, signal transduction, chromatin organization, DNA repair, and cytoskeletal organization (Figure 3F). These results suggest that, whereas ILS-associated loci are broadly distributed among housekeeping functions, introgression-associated loci are preferentially associated with regulatory and developmental processes that may contribute to phenotypic evolution.

Within the introgression-dominated node (node N7), discordant genes were further classified into V2- and V3-type according to the dominant discordant genealogy. Overall, V3-type genes substantially outnumbered V2-type genes, indicating that the EKIS-PGON-like discordant topology represented the predominant introgression signal across the genome (Figure 3E). Interestingly, comparison of exon- and intron-defined gene sets revealed strikingly different patterns between the two genealogical classes. Among V2-type genes, only four genes were shared between exon- and intron-based analyses, despite the two datasets containing similar numbers of genes (80 and 86 genes, respectively). In contrast, V3-type genes exhibited a much greater overlap, with 305 genes shared between exon- and intron-derived datasets (637 and 695 genes, respectively) (Figure 3E). These results indicate that exon and intron regions frequently retain distinct genealogical histories among V2-type loci, whereas V3-type loci tend to exhibit more consistent genealogical signals across coding and non-coding regions.

Within the introgression-dominated node (node N7), expression patterns of V2-type genes were compared between EKIS and PMAC. Overall, most V2-type genes exhibited divergent expression profiles between the two species (Supplementary Fig 15), consistent with previous observations that introgressed genomic regions frequently display heterogeneous transcriptional outcomes(56, 57). Nevertheless, a subset of genes retained highly similar tissue-specific expression patterns. Among these, *hpxa*, a liver-enriched gene involved in heme binding and oxygen metabolism, showed remarkably conserved expression across corresponding tissues in both species (Supplementary Table 11). Gene tree analysis further demonstrated that *hpxa* supported an EKIS– PMAC-like discordant topology rather than the species phylogeny (Figure 3D), suggesting that this locus represents a candidate introgressed gene that has retained conserved expression following introgression(58, 59). Similarly, *gng3*, a component of heterotrimeric G protein signaling, and *twf1a*, an actin-binding protein, also exhibited highly conserved tissue-specific expression patterns, with predominant expression in the brain and other corresponding tissues of both species (Supplementary Fig 15). Together, these examples indicate that although transcriptional divergence predominates among introgressed loci, a subset of introgression-associated genes has retained conserved regulatory programs following introgression, potentially preserving ancestral physiological or developmental functions.

### Plateau uplift and elevational gradients

To investigate how environmental gradients and regional geological processes have shaped molecular evolutionary dynamics in Sisoridae, we quantified genome-wide evolutionary rates and examined their variation across species, time, and genomic compartments. Ancestral state reconstruction of elevational distribution revealed a pronounced shift of Sisoridae lineages from low-to high-elevation habitats through time (Figure 4A). This transition was accompanied by substantial heterogeneity in genome-wide evolutionary rates among extant species. Boxplot comparisons of lineage-specific substitution rates showed marked variation across taxa (Figure 4B), indicating that molecular evolutionary tempo is not uniform across the phylogeny.

**Figure 4.**
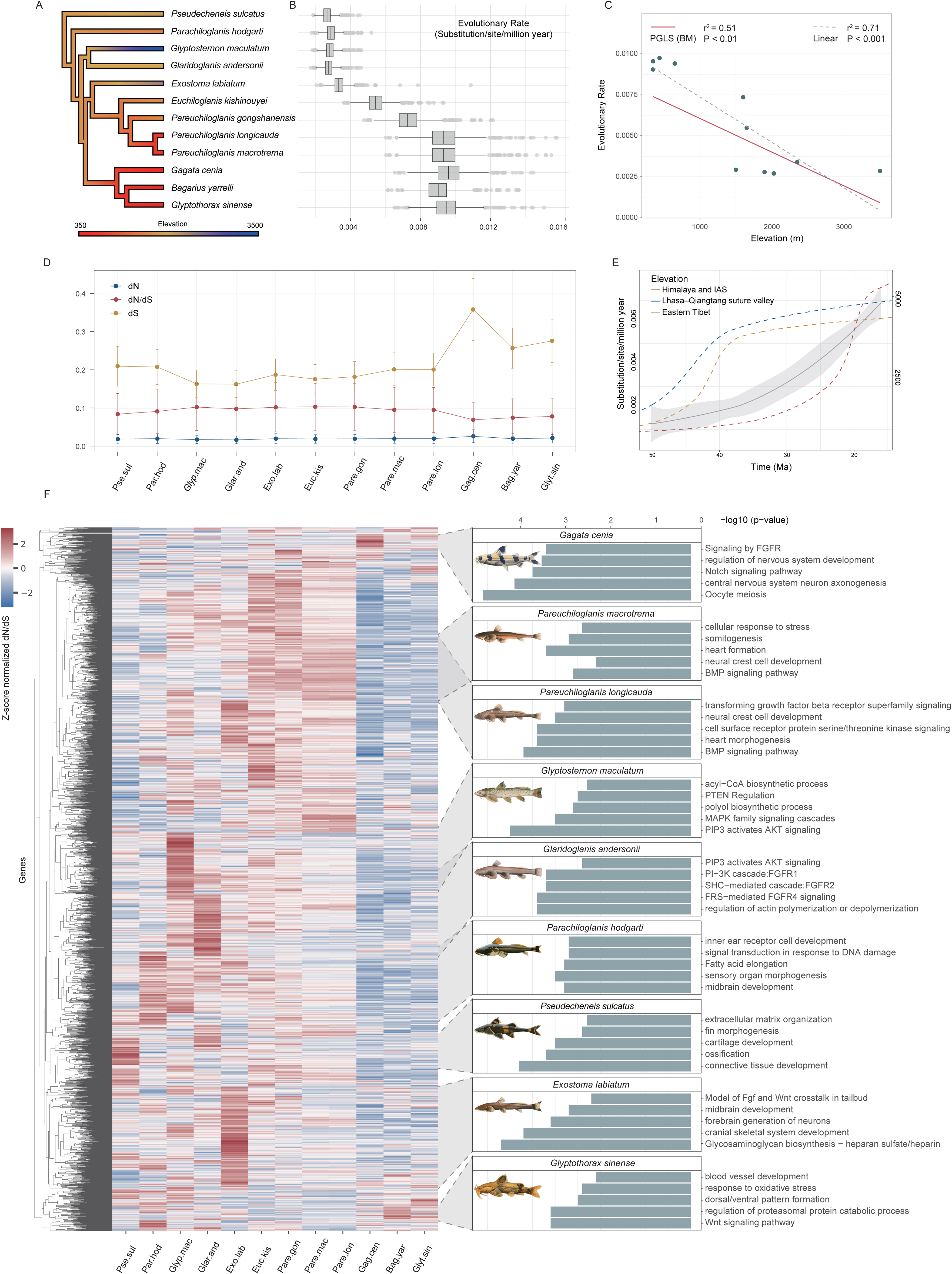
Evolutionary rate dynamics in Sisoridae in relation to the uplift of the Qinghai-Tibet Plateau and elevation gradients. **A.** Ancestral state reconstruction of elevation distributions across Sisoridae species. **B.** Boxplots of genome-wide evolutionary rates across species. **C.** Correlation between genome-wide evolutionary rates and elevation, where the red solid line and gray dashed line represent fits from phylogenetic generalized least squares (PGLS, BM model) and linear regression, respectively. **D.** Distribution patterns of synonymous substitution rates (dS), nonsynonymous substitution rates (dN), and their ratio (dN/dS) across protein-coding genes in Sisoridae species. **E.** Historical evolutionary rate dynamics of Sisoridae were reconstructed (gray solid line indicates the estimated values, with shaded areas representing confidence intervals) and compared with paleo-elevation changes across the Himalaya–Tibetan Plateau region. Red, blue, and yellow dashed lines represent the uplift histories of the Himalaya and Indo-Asian Suture (IAS), the Lhasa-Qiangtang suture zone, and the eastern Tibetan Plateau, respectively. **F.** Distribution of gene-level dN/dS Z-score values across species, highlighting lineage-specific patterns of relative evolutionary rate acceleration and functional enrichment of genes with extreme Z-scores.

To examine ecological correlates of this variation, we tested the relationship between genome-wide substitution rates and elevational distribution using both phylogenetic generalized least squares (PGLS; Brownian motion model) and standard linear regression. Both approaches consistently recovered a significant negative correlation between evolutionary rate and elevation (Figure 4C), with higher-elevation lineages exhibiting lower genome-wide substitution rates.

To place these patterns in a temporal context, we reconstructed evolutionary rate changes along the Sisoridae phylogeny using internal nodes and compared them with major phases of Qinghai-Tibet Plateau uplift history. The reconstructed rate results showed substantial temporal variation, with pronounced shifts occurring during the early diversification of the group (Figure 4E). When overlaid with regional tectonic events, including the India-Asia collision system (Himalaya-IAS), the Lhasa-Qiangtang suture zone activity, and the uplift of eastern Tibet, changes in evolutionary rates broadly coincided with major intervals of orogenic activity (Figure 4E), suggesting temporal concordance between lineage diversification dynamics and regional geological evolution(60).

Decomposition of substitution rates into synonymous (dS) and nonsynonymous (dN) components further revealed systematic differences among elevational groups. High-elevation species generally showed elevated dN/dS ratios, whereas low-elevation species exhibited higher dS values (Figure 4D), which is consistent with our earlier findings(41, 61).

These patterns indicate that variation in genome-wide rates reflects differential contributions of neutral and nonsynonymous substitutions across ecological gradients, rather than uniform shifts across all substitution classes. At the gene level, Z-score-normalized dN/dS matrices revealed strong heterogeneity in lineage-specific patterns of rate acceleration (Figure 4F). Different species showed distinct sets of rapidly evolving genes with limited overlap in composition. Functional enrichment analyses indicated that these lineage-specific gene sets were associated with different biological processes. For example, *Glyp. maculatum* and related lineages were enriched in pathways related to PI3K-AKT signaling and energy metabolism, whereas *Glyt. sinense* showed enrichment in genes associated with oxidative stress response. These patterns suggest that lineage-specific molecular rate shifts are accompanied by functional divergence in pathways relevant to ecological adaptation.

### Candidate genomic changes associated with torrent-related traits in Sisoridae

Members of Sisoridae inhabit fast-flowing mountain streams and display characteristic torrent-adapted morphologies, including dorsoventrally flattened bodies, enlarged paired fins, and specialized adhesive structures. Certain species additionally possess an accessory liver, which appears to be largely similar to the primary liver in its physiological and biochemical characteristics(62, 63). To explore the molecular basis of these traits, we analyzed lineage-specific positive selection, accelerated evolution, convergent substitutions, and variation in conserved non-coding elements across 12 species.

Based on major morphological features, species were grouped into distinct phenotypic categories, and genes showing positive selection or convergent evolution were identified for each group (Figure 5A). Although different morphological groups exhibited partially distinct functional enrichment profiles, the associated genes repeatedly converged on biological processes related to developmental regulation, skeletal and muscular system formation, and environmental response. These results suggest that morphological divergence in Sisoridae is shaped by changes across multiple functional systems rather than by alterations in a single genetic pathway (Supplementary Table 19, Supplementary Table 22).

**Figure 5.**
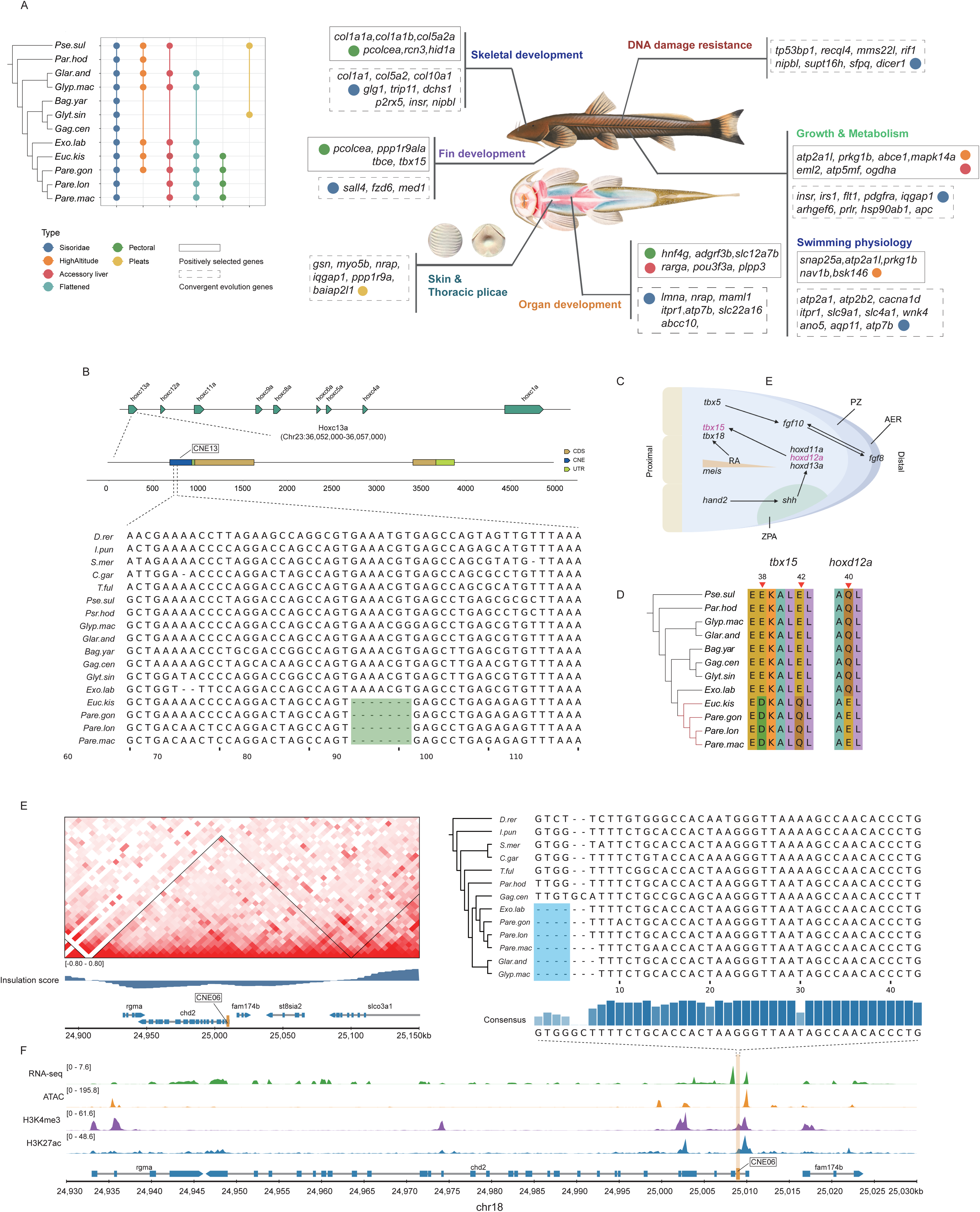
Candidate genomic changes associated with torrent-related traits in Sisoridae. **A.** Genes with unique evolutionary signals in Sisoridae species and their putative adaptive function. Top left: grouping of Sisoridae species based on distinct morphological traits. **B.** A differentially conserved non-coding element (dCNE) was identified in pectoral-fin-specialized lineages, located in the promoter region of the *hoxc13a* gene. **C.** Proposed roles of *tbx15* and *hoxd12a* in signaling pathways during pectoral fin bud development. **D.** Positively selected genes identified in pectoral-fin-specialized lineages and their amino acid substitution patterns. **E.** dCNEs within the same topologically associated domain (TAD) as *rgma* and *chd2* in flattened lineages. **F.** Genome tracks showing RNA-seq signal, H3K27ac and H3K4me3 histone modifications, and ATAC-seq chromatin accessibility in the genomic region surrounding the dCNE.

In pectoral fin-specialized lineages, we identified a divergent conserved non-coding element (dCNE) located in the promoter region of *hoxc13a* (Figure 5B). Sequence comparisons revealed lineage-specific insertion-deletion (indel) variation within this element, suggesting modification of local cis-regulatory structure. Transcription factor binding site predictions further indicated shifts in potential regulatory factor composition associated with these indels (Supplementary Table 24). In parallel, the fin-development gene *tbx15* showed signatures of positive selection in fin-specialized lineages, whereas *hoxd12a*, a gene previously implicated in the evolution of enlarged pectoral fins in Tibetan catfishes, exhibited lineage-specific convergent amino acid substitutions shared among fin-specialized taxa (Figure 5C,D)(41). Gene *tbx15* has been implicated in skeletal and mesenchymal development, whereas *hoxd12a* is involved in distal fin patterning and appendage morphogenesis(64–66). These substitutions are predicted to affect protein structure or functional domains, suggesting potential impacts on developmental signaling or cytoskeletal processes involved in fin morphogenesis.

For dorsoventrally flattened lineages, we identified a candidate conserved non-coding element with sequence variation near rgma and chd2 (Figure 5E).(67, 68).At the orthologous zebrafish locus, published adult muscle Hi-C data reveal local chromatin contact structure, while ATAC-seq, H3K27ac, and H3K4me3 tracks show signal over the mapped element (Figure 5E,F). The combination of lineage-associated sequence variation, proximity to genes involved in tissue patterning and transcriptional regulation, and muscle chromatin features at the zebrafish ortholog prioritizes this region as a candidate regulatory locus for further investigation.

We further examined genomic features associated with liver-related traits. The nuclear receptor gene *hnf4g* showed signatures of positive selection in Sisoridae, with positively selected sites located outside the DNA-binding and ligand-binding domains, specifically within a disordered C-terminal region enriched in polar residues, followed by a Sisoridae-specific 12-amino-acid deletion (Supplementary Fig 16). As a key transcription factor involved in endodermal organ development, HNF4 family genes play essential roles in liver, intestinal, and pancreatic development, epithelial differentiation, as well as lipid and carbohydrate metabolism(59). The lineage-specific deletion occurs within the predicted intrinsically disordered region of HNF4γ, while the conserved DNA-binding and ligand-binding domains remain intact (Supplementary Fig 16), suggesting that adaptive evolution may have preferentially targeted regulatory regions rather than the conserved functional core of the protein. Collectively, these patterns suggest that modulation of HNF4-mediated transcriptional programs may have contributed to hepatic developmental remodeling in Sisoridae, potentially facilitating the evolution of accessory liver structures as extensions of the primary liver rather than fully independent organs.

## Discussion

### Phylogenetic signal and early radiation of Siluriformes

Our phylogenomic analyses reveal a deep bifurcation of Siluriformes into two supra-regional lineages broadly corresponding to Pan-Asia and Pan-Africa, both tracing their origins to the late Mesozoic-early Cenozoic transition (∼66 Ma). Early divergences within Pan-Asia separated the Silur-Clari lineage from the Big-Asia clade, followed by extensive range expansion. The dispersal of Big-Asia into the Oriental region is consistent with an Indian Plate stepping-stone scenario during its northward drift and collision with Eurasia(48). In contrast, Pan-Africa lineages exhibit complex transcontinental histories, including secondary dispersal into Asia, and our results show that Ictaluridae diverged at ∼50 Ma and began diversifying at ∼28.6 Ma. This timing is consistent with the reported dynamics of trans-Beringian dispersal from Eurasia to North America: dispersal increased under warm climates after ∼65 Ma, was severely reduced by the ∼35 Ma glaciation, rebounded, and peaked at ∼25 Ma. The diversification of Ictaluridae at ∼28.6 Ma falls within this peak dispersal window, supporting climate-facilitated colonization of North America via the Bering land bridge(69, 70). Marine-tolerant clades such as Ariidae and Plotosidae further facilitated repeated oceanic and coastal dispersal, contributing to a near-global distribution by the Neogene(71).

Across genomic partitions, phylogenetic inference is strongly rate-dependent. Fast-evolving loci and third-codon positions consistently recover a stable backbone topology, whereas more conserved datasets yield conflicting or weakly resolved relationships(72). This pattern reflects short internode intervals during early cladogenesis, where limited substitution accumulation reduces phylogenetic signal in slowly evolving genes. Rather than representing methodological inconsistency, rate dependence likely captures the biological signature of rapid early radiation(73).

Divergence time analyses further support a complex diversification history across the K-Pg boundary. In contrast to many vertebrate groups that showed sustained diversification increases and post-K–Pg radiative expansion, Siluriformes experienced a long-term decline in diversification rates throughout the Late Cretaceous, reaching a minimum near the K–Pg boundary before recovering during the Paleogene.(46, 74, 75). This pattern may indicate that major siluriform lineages had already originated prior to the K-Pg extinction event, with subsequent diversification occurring during post-extinction ecological recovery rather than through immediate ecological release. The relatively longer branches observed during the Paleogene instead suggest a period of sustained lineage divergence and accumulation of genetic differentiation following the K-Pg bottleneck, consistent with gradual ecological recovery rather than explosive radiation. This interpretation aligns with fossil evidence showing a long-term decline of actinopterygians since the Permian, and a post-K-Pg crash in actinopterygian abundance linked to lower food-web disruptions. Against this backdrop, freshwater catfishes did not undergo immediate explosive radiation; rather, their gradual Paleogene diversification matches the slow ecological recovery, consistent with the longer branches we observe(76, 77).

Our results support a three-phase macroevolutionary model for Siluriformes: an early burst of diversification in the Early Cretaceous, a prolonged deceleration culminating in a K-Pg bottleneck, and a Paleogene rebound accompanied by rapid cladogenesis and extensive biogeographic expansion. This integrated scenario reconciles molecular divergence patterns with paleogeographic reconstructions and highlights the joint roles of Earth system reorganization and rate-dependent phylogenetic signal in shaping the evolutionary history of catfishes.

### Genomic discordance and complex evolutionary history of Sisoridae

Within Sisoridae, phylogenies based on whole-genome alignment data reject the monophyly of glyptosternoid fishes by nesting *Gagata*, *Bagarius*, *Glyptothorax*, and their ancestral lineage inside the glyptosternoid clade, whereas discordant gene trees— likely accounting for the monophyly recovered in previous single-gene studies—reveal substantial topological inconsistency(5–8). Together with widespread gene tree discordance, both introgression and incomplete lineage sorting (ILS) contributed to diversification and speciation within the family, although their relative influences likely differed across evolutionary stages in Sisoridae. During the early phase of diversification (∼50–35 Ma), corresponding to initial phases of Qinghai–Tibet Plateau uplift, geographic barriers among lineages may have remained relatively weak, permitting secondary contact and extensive introgression among diverging taxa(69). In contrast, introgression signals in later-diverging lineages were largely restricted to closely related species with geographically overlapping distributions. Compared with introgression, signals attributable to ILS appear to be relatively limited throughout the diversification history of Sisoridae.

These two processes appear to leave distinct genomic signatures. ILS signals are typically fragmented and locus-specific, reflecting stochastic sorting of ancestral polymorphisms during rapid divergence, and their retention is also associated with selective pressure(78). In contrast, introgressed regions tend to be more continuous and persist across larger genomic segments, suggesting longer-term retention following secondary contact.

Interestingly, discordant topologies associated with introgression also exhibited marked asymmetry. Across both genome-wide sites and gene-level analyses, the EKIS-PGON-like topology (V3) was substantially more prevalent than the EKIS-PMAC-like topology (V2). Moreover, exon- and intron-defined V3-type genes showed considerable overlap, whereas V2-type genes identified from coding and non-coding regions were almost completely distinct. These contrasting patterns suggest that the two discordant genealogies may represent introgression histories with different evolutionary persistence. One possible explanation is that the V3 signal reflects relatively continuous introgressed haplotypes that have been retained over longer evolutionary timescales, whereas the V2 signal has been more extensively reshaped by subsequent recombination, selection, or additional episodes of gene flow, resulting in fragmented genealogical histories between coding and non-coding regions.

An additional, non-mutually exclusive explanation may relate to the evolutionary history of *Pareuchiloglanis gongshanensis*. Unlike the more widely distributed congeners, *P. gongshanensis* is geographically restricted to the upper Nujiang River. If ancestral introgression between the lineage leading to *P. gongshanensis* and *Euchiloglanis* occurred prior to geological isolation of this drainage, subsequent long-term isolation may have facilitated the preservation of ancestral introgressed haplotypes. In contrast, more widespread species such as *P. macrotrema* may have experienced repeated recombination, population turnover, and continued gene flow with related lineages, progressively eroding earlier introgression signatures and leaving only a relatively small subset of V2-type loci. Although this scenario remains speculative, it provides a plausible explanation for the pronounced asymmetry observed between the two discordant genealogies.

Such introgressed variation may represent an important source of genetic diversity, potentially contributing to phenotypic diversification and ecological differentiation in heterogeneous riverine environments(79). Although coalescent hidden Markov models (coalHMMs) have been widely applied to infer incomplete lineage sorting among closely related species, we further incorporated QuIBL to explicitly evaluate topological discordance across gene trees, enabling more robust discrimination between signals generated by ILS and those arising from introgression(78, 80, 81). Our results suggest that the combined effects of rapid divergence and episodic gene flow likely underlie the persistent phylogenetic uncertainty observed within Sisoridae.

### Genome evolution, demography, and environmental association

Transposable elements (TEs) are recognized as important drivers of genome evolution, and teleost fishes exhibit substantially greater TE diversity than most other vertebrate groups(82–84). Patterns of genome size and transposable element (TE) content differ markedly between major ecological groups within Sisoridae. Torrent-adapted taxa, particularly glyptosternine lineages, generally possess larger genomes (∼700–900 Mb) and substantially higher TE proportions (∼30–40%), whereas non-torrent taxa tend to exhibit more compact genomes (∼500–600 Mb) with lower TE content. In teleosts, genome size variation is strongly associated with TE abundance, and genome contraction is often accompanied by reductions in TE copy number, likely associated with purifying selection. Under reduced effective population size and relaxed selective constraints, TE proliferation is more readily tolerated, promoting genome enlargement(85, 86). These differences are consistent with reduced long-term effective population sizes (Ne) in fragmented montane river systems, where weakened purifying selection may facilitate the accumulation of TE insertions and contribute to genome expansion.

Molecular evolutionary rates also differ between ecological groups. Torrent-adapted glyptosternoid fishes exhibit relatively elevated genome-wide dN/dS ratios compared with non-torrent taxa, although all species retain values below 1, indicating the predominance of purifying selection. Elevated dN/dS in torrent-adapted species may reflect a combination of relaxed selective constraint and episodic positive selection associated with adaptation to extreme montane environments. This pattern mirrors observations in island species(87).

Importantly, these genomic signatures are consistent with the broader evolutionary history reconstructed in this study. Time-calibrated diversification analyses indicate that the major radiation of Sisoridae closely tracked successive phases of Qinghai-Tibet Plateau uplift, whereas ancestral-state reconstruction suggests repeated ecological transitions into high-elevation montane river systems during sisorid evolution(25, 60). Moreover, phylogenetically informed comparative analyses revealed a significant negative association between evolutionary rate and elevation, indicating that environmental transitions have influenced not only lineage diversification but also the tempo of genome evolution. Those results are consistent with a possible relationship between regional geological change and sisorid diversification(88, 89).

Although direct estimates of historical Ne remain limited, the combined genomic patterns suggest that demographic history has played an important role in shaping genome evolution within Sisoridae. Habitat fragmentation, reduced connectivity, and long-term population subdivision in mountain river systems may jointly influence both genome architecture and molecular evolutionary dynamics by reducing the efficacy of purifying selection while permitting localized adaptive evolution(90–92). For example, similar patterns of reduced purifying selection and an increased proportion of nonsynonymous mutations associated with habitat fragmentation have been documented in both frogs and mammals inhabiting montane regions(93, 94). These findings have important implications for species conservation: maintaining habitat connectivity is critical for preserving genetic diversity, mitigating the accumulation of deleterious mutations, and sustaining the long-term adaptive potential of fragmented populations.

### Heterogeneous genomic basis of adaptation in Sisoridae

Our analyses indicate that adaptation in Sisoridae has proceeded through multiple and partially independent genomic routes rather than through a single shared genetic mechanism (Figure 5). Signals of accelerated evolution, positive selection, and convergent amino acid substitution were detected across different lineages and species, yet the underlying candidate genes showed limited overlap (Figure 4F). Despite this heterogeneity at the gene level, many of these genes converge functionally on processes associated with environmental adaptation and morphological specialization, including skeletal and muscular development, fin morphogenesis, adhesion-related functions, developmental regulation, and metabolic or stress-response pathways. These patterns suggest that similar ecological pressures in torrent habitats may repeatedly target overlapping functional systems while utilizing distinct genetic substrates in different lineages(95, 96).

A notable example involves the evolution of accessory liver structures in Sisoridae. Within Sisoridae, the development of accessory liver structures has been reported to correlate with increasing dorsoventral body flattening, suggesting that their formation may represent a regulated developmental feature rather than a purely pathological condition(97). We identified a liver-development-related gene, *hnf4g*, exhibited a conserved amino acid replacement and lineage-specific deletion across all examined sisorid species, including both accessory-liver and non-accessory-liver taxa. Because the observed hnf4g changes are shared by taxa with and without accessory livers, they cannot explain the presence of that structure on their own. Functional experiments and trait-matched expression data would be needed to test whether hnf4g contributes to accessory-liver development(98–100).

These findings support a model in which the ecological and morphological diversification of Sisoridae arose through heterogeneous combinations of coding and regulatory evolution. Similar adaptive phenotypes may therefore emerge through different genomic trajectories acting on partially overlapping developmental and physiological systems.

## Method

### Taxon sampling

Resolving the evolutionary origin and phylogenetic placement of Sisoridae represents a longstanding challenge in Siluriformes systematics, primarily due to limited taxon sampling and insufficient molecular data. Previous phylogenetic studies of Siluriformes have been constrained by sparse marker sampling, typically relying on mitochondrial genomes or a small number of nuclear loci, resulting in insufficient resolution of higher-level relationships across the order. To address these issues, with a particular focus on clarifying the origin and diversification of Sisoridae while also improving phylogenetic resolution across Siluriformes, we constructed a comprehensive phylogenomic dataset based on large-scale genomic resources with broad taxonomic coverage. Our sampling framework was established following the taxonomic classifications of Betancur-R et al., Nelson et al., and the updated synthesis of Siluriformes systematics by Near & Thacker, in combination with the Catalog of Fishes(1, 2, 20). According to these taxonomic references, extant Siluriformes comprise 40 recognized families. Following this framework, we systematically surveyed all publicly available molecular resources for Siluriformes, including both genome assemblies and transcriptome datasets. Data quality was assessed using BUSCO, and only datasets with a completeness score exceeding 70% (complete BUSCOs) were retained for our analyses. Based on this framework, our dataset includes representatives from 28 families, spanning 107 genera and a total of 180 species, thereby providing extensive coverage across major siluriform lineages (Supplementary Table 1). Sampling was designed to encompass all three suborders within Siluriformes, including Loricarioidei (one family sampled), Diplomystoidei (three of six families sampled), and Siluroidei (24 of 33 families sampled). The selected taxa represent a wide geographic distribution, covering Eurasia, Southeast Asia, Africa, and the Americas, ensuring a globally representative dataset for the order. In addition to broad taxonomic coverage, this study substantially expands genomic resources for several key lineages. To specifically resolve relationships within Sisoridae, we generated chromosome-level genome assemblies for 7 species and scaffold-level assemblies for 2 species within this family and, together with publicly available datasets, reconstructed the phylogenetic relationships among 12 genera of East Asian sisorid catfishes for the first time. Furthermore, our sampling enhances representation of previously under-sampled families, including two species from Amblycipitidae, three from Amphiliidae, and one from Mochokidae.

### DNA extraction, library preparation and sequencing

Fresh muscle or liver tissues were collected from all newly sequenced species and preserved in 95% ethanol or liquid nitrogen prior to DNA extraction. Genomic DNA was extracted using a standard phenol-chloroform protocol with RNase A and proteinase K treatment. DNA quality and integrity were assessed by agarose gel electrophoresis, Qubit fluorometric quantification and NanoDrop spectrophotometry. High-molecular-weight (HMW) DNA (≥ 50 kb) was selected for long-read library construction.

### PacBio HiFi library preparation

PacBio HiFi libraries were constructed for *Parachiloglanis hodgarti*, *Glaridoglanis andersonii*, *Exostoma labiatum*, *Pareuchiloglanis longicauda*, *Pareuchiloglanis macrotrema*, *Gagata cenia*, *Amphilius grammatophorus*, *Phractura sp.*, *Belonoglanis tenuis* and *Synodontis eupterus*. Libraries were prepared using the SMRTbell protocol with size selection to enrich fragments >15 kb. Library quality and concentration were assessed prior to sequencing.

### PacBio CLR library preparation

For *Pareuchiloglanis gongshanensis*, a PacBio long-read library was constructed using the SMRTbell protocol with size selection for large DNA fragments. This approach generates continuous long reads without circular consensus sequencing.

### Oxford Nanopore library preparation

Oxford Nanopore libraries were prepared for *Pseudecheneis sulcatus* and *Glyptothorax sinense* using ligation-based protocols. HMW DNA was enriched for long fragments, and libraries were constructed following standard procedures. Library quality was assessed prior to sequencing.

### Illumina library preparation

Illumina paired-end libraries were constructed for *Xiurenbagrus gigas* and *Liobagrus anguillicauda*. Genomic DNA was fragmented to an average insert size of ∼350 bp, followed by adapter ligation and PCR amplification. Library quality was assessed prior to sequencing.

### Hi-C library preparation

Hi-C libraries were constructed for *Parachiloglanis hodgarti*, *Glaridoglanis andersonii*, *Exostoma labiatum*, *Pareuchiloglanis longicauda*, *Pareuchiloglanis macrotrema*, *Pareuchiloglanis gongshanensis*, *Gagata cenia*, *Amphilius grammatophorus*, *Phractura sp.* and *Belonoglanis tenuis.* Cross-linked chromatin was digested with HindIII, followed by end repair, biotin labeling, proximity ligation and library construction according to standard Hi-C protocols.

### Sequencing

PacBio HiFi libraries were sequenced on the Sequel II platform to generate high-fidelity long reads. The PacBio CLR library was sequenced to generate continuous long reads. Oxford Nanopore libraries were sequenced on the PromethION platform. Illumina libraries were sequenced on the NovaSeq 6000 platform to generate 150 bp paired-end reads.

### Public data collection

In addition to the newly generated datasets, we incorporated publicly available genomic and transcriptomic resources from Siluriformes species. These included assembled genomes, transcriptome assemblies and whole-genome resequencing data generated using Illumina sequencing. The collected data span a range of assembly qualities, including chromosome-level genomes, draft assemblies, transcriptome-derived sequences and raw sequencing reads. All data were obtained from NCBI (https://www.ncbi.nlm.nih.gov/) and other relevant repositories. Detailed information on species and data sources is provided in Supplementary Table 1.

### RNA extraction and sequencing

To provide transcriptomic evidence for genome annotation, RNA was extracted from multiple tissues of representative Siluriformes species. For *Parachiloglanis hodgarti*, *Glaridoglanis andersonii*, *Exostoma labiatum* and *Gagata cenia*, RNA was extracted from pooled tissues (brain, fin, gill, heart, liver, muscle and spleen). For *Pareuchiloglanis longicauda*, *Pareuchiloglanis macrotrema* and *Euchiloglanis kishinouyei*, RNA was obtained from brain, fin, gill, heart, liver, muscle and spleen tissues. Total RNA was extracted using TRIzol reagent (Invitrogen) and assessed for quality using standard electrophoresis and spectrophotometric quantification. RNA-seq libraries were constructed using the NEBNext Ultra RNA Library Prep Kit for Illumina (NEB, USA) following standard protocols, and sequenced on the Illumina NovaSeq platform to generate 150 bp paired-end reads.

### Genome assembly and Hi-C scaffolding

#### *De novo* genome assembly

For species sequenced using PacBio HiFi reads, including *Parachiloglanis hodgarti*, *Glaridoglanis andersonii*, *Exostoma labiatum*, *Pareuchiloglanis longicauda*, *Pareuchiloglanis macrotrema*, *Gagata cenia*, *Amphilius grammatophorus*, *Phractura sp.*, *Belonoglanis tenuis* and *Synodontis eupterus*, *de novo* genome assemblies were generated using hifiasm with default parameters(101). HiFi reads were directly assembled into high-quality primary contigs. For *Pareuchiloglanis gongshanensis*, sequenced using PacBio Sequel (CLR) data, de novo assembly was performed using Canu, followed by downstream scaffolding. For *Pseudecheneis sulcatus* and *Glyptothorax sinense*, sequenced using Oxford Nanopore long reads, de novo genome assemblies were generated using Flye with default parameters(102). For *Xiurenbagrus gigas* and *Liobagrus anguillicauda*, genome assemblies were generated from short-read data using SOAPdenovo2 based on a de Bruijn graph approach(103). For *Chaca chaca*, raw Illumina sequencing data were downloaded from NCBI (ERR14369883) and assembled using Flye with default parameters. Assembly quality was evaluated using standard metrics, including total assembly size and contig N50. Genome completeness was assessed using BUSCO with an appropriate lineage dataset(104).

### Hi-C scaffolding

Hi-C-based scaffolding was performed for *Parachiloglanis hodgarti*, *Glaridoglanis andersonii*, *Exostoma labiatum*, *Pareuchiloglanis longicauda*, *Pareuchiloglanis macrotrema*, *Pareuchiloglanis gongshanensis*, *Gagata cenia*, *Amphilius grammatophorus*, *Phractura sp.* and *Belonoglanis tenuis*. Raw Hi-C reads were first processed using fastp to remove adapter sequences and low-quality bases(105). Clean reads were initially mapped to the draft genome assemblies using Bowtie2 (v2.5.5)(106). Reads spanning ligation junctions that failed to map were further processed by restriction site-based trimming, followed by re-mapping to the reference genome. After alignment, valid Hi-C read pairs were identified by filtering out unmapped reads, PCR duplicates, self-ligated fragments and other invalid interaction pairs. Based on the filtered valid pairs, contact matrices were constructed at multiple resolutions to represent chromatin interaction frequencies across the genome. Chromosome-scale scaffolding was performed using 3D-DNA with default parameters, followed by manual curation using Juicebox (https://github.com/aidenlab/Juicebox) to correct misjoins and improve scaffold accuracy(107).

### *De novo* transcriptome assembly and quality assessment

Raw paired-end RNA-seq reads were first assessed for quality using FastQC (https://github.com/s-andrews/FastQC). Low-quality reads, adapter contamination and ambiguous bases were removed to obtain clean reads for downstream analyses. *De novo* transcriptome assemblies were generated using Trinity with default parameters(108).

To reduce redundancy in the assembled transcripts, highly similar sequences were clustered using CD-HIT(109). To further improve transcriptome representation at the gene level, assembled transcripts were hierarchically clustered based on read mapping and expression patterns using Corset, and the longest transcript from each cluster was retained as a representative unigene set(110). The completeness of each transcriptome assembly was evaluated using BUSCO with an appropriate lineage dataset. Transcriptome assemblies with BUSCO completeness lower than 40% were excluded from subsequent analyses.

### Phylogenetic inference for Siluriformes

A total of 180 species representing 28 families of Siluriformes were included based on available genome and transcriptome datasets. Single-copy orthologous genes were identified using BUSCO with a species occupancy threshold of 85%–90%. For species represented by transcriptome data, duplicated BUSCO genes were retained to supplement the ortholog set. In total, 1,125–1,896 BUSCO genes were selected for downstream analyses. For the 1,896 high-coverage BUSCO genes, six datasets were constructed: (1) full codon alignments, (2) first and second codon positions (P1+P2), (3) third codon positions (P3), (4) amino acid sequences, (5) the fastest-evolving 50% of genes, and (6) the most conserved 50% of genes. Gene evolutionary rates were obtained from the OrthoDB database, and genes were ranked accordingly and divided into the top 50% (fast-evolving) and bottom 50% (conserved) groups. Each dataset was aligned and concatenated into supermatrices for phylogenetic reconstruction under a maximum likelihood (ML) approach.

Phylogenetic analyses were conducted using IQ-TREE2(111). For nucleotide (CDS) datasets, trees were inferred under the GTR+G+gap model, whereas amino acid datasets were analysed under the JTT+G+gap model. Branch support was assessed using ultrafast bootstrap with 1,000 replicates (-B 1000) and SH-like approximate likelihood ratio tests (-alrt 1000), with branch length optimization using nearest-neighbour interchange (-bnni). To account for gene tree heterogeneity, individual gene trees were reconstructed for all BUSCO loci and used to infer a coalescent-based species tree with ASTRAL under the multi-species coalescent (MSC) approach(112).

To root the phylogeny, multiple non-siluriform teleost species were included as outgroups, including *Clupea harengus*, *Scleropages formosus*, *Danio rerio*, *Electrophorus electricus*, and *Astyanax mexicanus*. For visualization purposes, only *Danio rerio* was retained in the final tree, while the remaining outgroup lineages were pruned to improve clarity.

### Phylogenetic discordance analysis

To assess phylogenetic incongruence, gene trees were reconstructed for all 1,125-1,896 BUSCO loci using IQ-TREE2 with automatic model selection and 1,000 ultrafast bootstrap replicates. Gene tree discordance was quantified using DiscoVista by comparing individual gene trees with the reference species tree(113). The proportions of concordant, conflicting and uninformative gene trees were calculated for each focal branch, allowing visualization of gene tree conflict and providing insights into potential sources of discordance, including incomplete lineage sorting, introgression and systematic bias.

### Divergence time estimation and fossil calibration

Divergence times were estimated using MCMCTree based on the species tree inferred from the concatenated BUSCO dataset (all codon positions), with the same alignment used for both phylogenetic reconstruction and molecular dating(114). A total of 12 calibration points were applied, including fossil, secondary, and geological constraints distributed across Teleostei, Otomorpha, Siluriformes, and multiple internal nodes within Siluroidei. Deep nodes were constrained using well-established fossil records (Teleostei: 215-252.7 Ma; Otomorpha: 150.9-228.4 Ma; Siluroidei: 65.5-146.5 Ma), while additional minimum or bounded constraints were applied to family- and species-level divergences. Detailed information on all priors and calibration points is provided in Supplementary Table 2. Uniform priors were applied to most nodes, with a gamma prior used for the Ariidae + Anchariidae split. A relaxed molecular clock model was implemented to accommodate rate variation among lineages.

### Rate-through-time analysis

We quantified diversification dynamics using Bayesian Analysis of Macroevolutionary Mixtures (BAMM, v2.5). A time-calibrated phylogeny comprising 181 (include 1 outgroup species *Danio rerio*) species was used as the input tree.We used a species level tree and diversification rate analysis was conducted under a speciation-extinction model with Markov chain Monte Carlo (MCMC) sampling for 100 million generations. The temperature increment parameter was set to 0.01, and event data were sampled every 1,000 generations. The first 20% of samples were discarded as burn-in. Diversification rate-through-time curves were estimated and visualized in BAMMtools(115).

### Genome annotation

#### Repeat annotation

Repetitive elements in all genomes were annotated using a combination of *de novo* and homology-based approaches. For de novo identification, RepeatModeler (v1.0.5) (https://github.com/Dfam-consortium/RepeatModeler) was used to construct a species-specific transposable element library. The resulting library, together with known repeats from Repbase, was used as input for RepeatMasker (v4.0.6) to annotate genomic repeats(116). Homology-based repeat identification was also performed using RepeatMasker and RepeatProteinMask against the Repbase database. Tandem repeats were further identified using TRF (v4.0.4)(117). The overall composition and distribution of repeat elements across genomes were summarized for comparative analyses (Supplementary Table 4).

### Protein-coding gene prediction

Protein-coding genes were annotated using a combined strategy integrating *ab initio* prediction, homology-based inference, and transcriptome evidence where available. For *ab initio* prediction, gene models were generated using AUGUSTUS, GlimmerHMM and SNAP with default or species-trained parameters(118–120). For homology-based prediction, protein sequences from representative vertebrate genomes were aligned to repeat-masked genomes using TBLASTN (E-value < 1e−5), followed by gene structure refinement using Genewise(121, 122). For transcriptome-based prediction, RNA-seq reads were mapped to the genome using STAR, and transcripts were assembled using StringTie(123, 124). Transcript structures were further refined by PASA (https://github.com/PASApipeline/PASApipeline/tree/master) to improve exon–intron boundaries and alternative isoform annotation. All gene predictions were integrated using EvidenceModeler (EVM) to generate a non-redundant consensus gene set, with transcript evidence assigned higher weights than ab initio predictions(125). Final gene models were filtered to remove potential transposable element-derived sequences. (Supplementary Table 4)

### Functional annotation

Protein sequences were functionally annotated by similarity searches against the SwissProt, TrEMBL and NCBI NR databases using BLASTP (E-value < 1e−5)(126). Protein domains were identified using InterProScan against Pfam, SMART, PROSITE and related databases. Gene Ontology (GO) terms were assigned based on InterPro and SwissProt mappings. KEGG pathway annotations were performed using KOBAS(127–129).

### Whole-genome alignment

Whole-genome alignments (WGAs) were generated for 12 representative species of Sisoridae, including *Pseudecheneis sulcatus*, *Parachiloglanis hodgarti*, *Glyptosternon maculatum*, *Glaridoglanis andersonii*, *Exostoma labiatum*, *Euchiloglanis kishinouyei*, *Pareuchiloglanis gongshanensis*, *Pareuchiloglanis longicauda*, *Pareuchiloglanis macrotrema*, *Gagata cenia*, *Bagarius yarrelli*, and *Glyptothorax sinense*. These taxa were selected to represent the major lineages within Sisoridae, enabling detailed investigation of their phylogenetic relationships and potential sources of phylogenetic discordance. Pairwise genome alignments were first generated against the reference genome of Ictalurus punctatus, and subsequently combined into multiple alignments using the LASTZ + MULTIZ pipeline(130). The resulting alignments were processed in MAF format. Alignment blocks shorter than 100 bp or lacking any of the focal species were removed, and the remaining blocks were sorted and partitioned by chromosome to facilitate downstream analyses. To improve alignment quality and retain phylogenetically informative regions, we applied a series of filtering criteria. Alignment windows were excluded if (i) any species contained more than 30% gap sites, (ii) sequences from two or more species were identical across the window, or (iii) more than 70% of sites were identical across all species. After filtering, a total of 138,150 WGA blocks were retained from an initial alignment length of ∼213 Mb.

### Phylogenetic inference for Sisoridae

Each retained WGA block was converted into phylogenetic format and used for gene tree inference under a maximum likelihood (ML) framework implemented in IQ-TREE2. The best-fitting substitution model for each locus was selected using ModelFinder, and node support was assessed using ultrafast bootstrap approximation (UFBoot). Gene trees failing the χ² test for compositional heterogeneity were excluded, resulting in 120,858 high-quality gene trees for downstream analyses. To evaluate phylogenetic discordance across the genome, we quantified topological variation among gene trees and visualized conflicting signals using DiscoVista for every node. In addition, a coalescent-based species tree was reconstructed using ASTRAL based on the filtered gene tree set, allowing us to assess the extent of incomplete lineage sorting (ILS) and potential introgression among major sisorid lineages.

### QuIBL analysis

To disentangle the relative contributions of incomplete lineage sorting (ILS) and introgression to phylogenetic discordance, we performed QuIBL analyses based on locus-specific trees inferred from whole-genome alignment (WGA) data(131). Whole-genome alignments were partitioned by chromosome, and local phylogenetic trees were reconstructed independently for each chromosome. In total, 120,858 locus trees across all chromosomes were generated and used as the initial input. Trees containing anomalous branch lengths were excluded (min_branch_length = 1 × 10-⁶), yielding a final dataset of 119,251 locus trees. QuIBL analyses were performed independently for each chromosome using the corresponding filtered tree set, and the resulting outputs were subsequently pooled and summarized by species triplets for ΔBIC calculation.

QuIBL analyses were conducted for each internal node of the Sisoridae species tree by constructing node-specific triplet configurations. For each focal node, all possible combinations of three taxa (or taxon groups) consistent with the tested phylogenetic relationship were enumerated. Specifically, eight focal nodes were evaluated: node1 (PSUL-PHOD-X); node2 (PHOD-GAND-X and PHOD-GMAC-X); node3 (group1_group2_group3; group1: GMAC, GAND; group2: EKIS, ELAB, PGON, PLON, PMAC; group3: GSIN, GCEN, BYAR); node4 (ELAB–group2–group3, group2: EKIS, PGON, PLON, PMAC, group3: GSIN, GCEN, BYAR); node5 (GSIN-GCEN-BYAR); node6 (ELAB-EKIS-group2, group2: PGON, PLON, PMAC); node7 (EKIS-PGON-X), and node8 (PGON-PLON-PMAC). For each triplet, loci trees were partitioned into three topological subsets corresponding to the alternative resolutions of the three taxa, with the topology consistent with the species tree treated as concordant and the remaining two treated as discordant. For visualization of introgression intensity, the estimated total introgression proportion (totalIntroProp) from individual QuIBL analyses was subsequently grouped according to the corresponding lineage combinations within each focal node and summarized across all eligible triplets (Supplementary Fig 14).

For each discordant subset, QuIBL evaluated whether the distribution of internal branch lengths was better explained by a model assuming ILS alone (single exponential distribution) or a model incorporating both ILS and introgression (mixture model). Model fit was assessed using the Bayesian Information Criterion (BIC). The difference in BIC values (ΔBIC = BIC_mixture-BIC_ILS) was used to distinguish between scenarios: ΔBIC > 10 supports ILS as the primary driver of discordance, whereas ΔBIC < −10 supports a mixed model involving introgression; values between −10 and 10 were considered inconclusive. In addition, QuIBL estimated the expected distributions of internal branch lengths under each model, allowing comparison between theoretical and observed patterns of phylogenetic discordance. For each internal node, QuIBL results from all discordant triplets were subsequently summarized to characterize the predominant evolutionary process. Nodes in which fewer than 50% of discordant triplets produced significant model support (|ΔBIC| > 10) were classified as inconclusive. For the remaining nodes, the relative proportions of significant triplets supporting the ILS-only model (BIC1) or the introgression model (BIC2) were compared, and each node was classified as ILS-dominated or introgression-dominated according to the majority-supported model (Supplementary Table 7).

### AutoCoalHMM analysis

To characterize local genealogical variation and quantify phylogenetic discordance across the genome, we applied AutoCoalHMM to whole-genome alignments (WGA)(132). Based on the WGA analyses described above, approximately 213 Mb of high-confidence aligned genomic sequences shared among all analyzed taxa were retained after filtering and used as input for AutoCoalHMM. Guided by the QuIBL results, two representative internal nodes were selected for detailed investigation, corresponding to nodes predominantly shaped by incomplete lineage sorting (ILS; node5) and introgression (node7), respectively. For each node, a fixed four-taxon configuration was defined according to the inferred species tree. Specifically, node5 was represented by the topology (((BYAR, GSIN), GCEN), IPUN), and node7 by (((PMAC, PGON), EKIS), IPUN). AutoCoalHMM was applied to the corresponding WGA data to infer local genealogical states along the genome. For each genomic position, posterior probabilities of four hidden states (V0-V3) were obtained, representing alternative genealogical topologies, including those concordant with the species tree and those reflecting discordant evolutionary histories.

Site-wise genealogical states inferred by AutoCoalHMM were mapped onto the Ictalurus punctatus reference genome and annotated according to gene models. Genomic positions were classified as exonic, intronic, or intergenic based on overlap with annotated gene features. For each position, the hidden state with the highest posterior probability was assigned as the genealogical state when the maximum posterior probability exceeded 0.35; otherwise, the site was considered ambiguous and excluded from downstream analyses. States V0 and V1 were regarded as concordant genealogical states, whereas V2 and V3 were regarded as discordant genealogical states.

To summarize genome-wide patterns, the proportions of V0-V3 states were quantified across exonic, intronic, and intergenic regions for each node. Gene-level analyses were performed using all genic sites, including both exon and intron regions. Informative sites were defined as sites assigned to V0, V1, V2, or V3 after excluding ambiguous sites. Discordant sites were defined as informative sites assigned to either V2 or V3. For each gene, the discordance ratio was calculated as the proportion of discordant sites among all informative sites:

Discordance ratio = (V2 + V3) / (V0 + V1 + V2 + V3)

To ensure robust gene-level estimates and minimize the influence of genes represented by only a small number of aligned positions, minimum informative-site thresholds were applied. For analyses based on all genic sites or intronic sites, only genes containing at least 100 informative sites were retained. For exon-specific analyses, a relaxed threshold of at least 50 informative sites was used because coding regions are generally shorter than intronic regions. These filtering criteria reduce the impact of stochastic variation associated with sparse alignment coverage and provide more reliable estimates of local genealogical composition.

Genes with a discordance ratio greater than 0.5 were considered to exhibit elevated genealogical discordance. Based on the node-specific evolutionary context inferred from QuIBL analyses, discordant genes identified at node5 were interpreted as primarily associated with incomplete lineage sorting, whereas discordant genes identified at node7 were considered candidates associated with introgression. These gene sets, defined using all genic sites, were subsequently used for comparisons of functional annotations and gene ontology categories between ILS- and introgression-associated genes.

For node7, corresponding to the four-taxon configuration (((PMAC, PGON), EKIS), IPUN), introgression-associated genes were further partitioned according to the predominant discordant genealogical state. Specifically, genes containing more V2 than V3 sites were classified as V2-type genes, whereas genes containing more V3 than V2 sites were classified as V3-type genes. Based on the genealogical interpretation of the hidden states, V2-type genes support an EKIS-PMAC-like discordant topology (((PMAC, EKIS), PGON), IPUN), whereas V3-type genes support an EKIS-PGON-like discordant topology (((PGON, EKIS), PMAC), IPUN).

To investigate the potential functional consequences of introgression, exon- and intron-specific discordant signals were analyzed separately for node7. Genes showing predominant V2 signals in exonic regions were examined for coding-sequence variation and potential protein-level functional changes, whereas genes showing predominant V2 signals in intronic regions were further evaluated for expression differences between PMAC and EKIS. This framework allowed the potential coding and regulatory consequences of introgression to be assessed independently.

Functional enrichment analyses of genes affected by incomplete lineage sorting (ILS) or introgression were performed using Gene Ontology (GO) annotation in the R package clusterProfiler to assess potential functional enrichment associated with genealogical discordance.

### Inference of ancestral linkage groups (ALGs)

Ancestral linkage groups (ALGs) in Sisoridae were inferred using the software odp (v0.3.066) based on chromosome-level genome assemblies(133). The analysis included eight representative species: *Pareuchiloglanis longicauda, Pareuchiloglanis macrotrema, Glyptosternon maculatum, Euchiloglanis kishinouyei, Gagata cenia, Parachiloglanis hodgarti, Glaridoglanis andersonii,* and *Exostoma labiatum*. To identify conserved orthologous relationships across species, we first performed an N-way reciprocal best hit (RBH) analysis using odp_nway_rbh with num_permutations set to 1,000,000. Pairwise .groupby outputs were subsequently converted into RBH format using odp_groupby_to_rbh, and all RBH results were merged across species using odp_rbh_merge to generate a unified orthology file, which served as the basis for ALG inference. Based on the merged RBH dataset, we constructed genome-wide alignment blocks using odp_rbh_to_alignments, which were then used as input for the main odp program to infer ancestral linkage relationships and reconstruct ALGs across Sisoridae genomes. For visualization, ribbon plots representing chromosomal correspondences and rearrangements among species were generated using rbh_to_ribbon, based on the inferred RBH and ALG relationships.

### Chromosomal synteny and 3D genome organization analyses

To investigate chromosomal structural variation in Sisoridae, we performed genome-wide synteny analyses using the JCVI utility libraries. Protein-coding genes from each genome were compared in an all-versus-all manner using BLASTP, and syntenic blocks were identified using the MCscan module implemented in JCVI with default parameters(134). The resulting collinearity relationships were used to identify large-scale chromosomal rearrangements, including fusion and fission events across species. To characterize genomic features associated with chromosomal rearrangements, genome-wide distributions of gene density, transposable elements (TEs), and tandem repeats (TRs) were calculated using a sliding window approach along each chromosome. TE and TR annotations were obtained as described above, and gene coordinates were derived from genome annotation results.

For three-dimensional genome organization, Hi-C data were processed using the HiCExplorer suite. Raw Hi-C reads were aligned to the reference genome and filtered to remove low-quality, unmapped, and duplicate reads(135). Contact matrices were constructed and stored in .cool format using cooler, which facilitates efficient storage and manipulation of Hi-C interaction data. Matrices were subsequently normalized (ICE normalization) to correct for biases in sequencing depth and genomic features. Topologically associating domains (TADs) were identified using hicFindTADs based on the insulation score (IS) method. Briefly, insulation scores were calculated across the genome using sliding windows, and TAD boundaries were defined as local minima in the insulation profile. Hi-C contact maps and insulation score tracks were used to visualize chromatin interaction patterns and domain organization.

### Demographic history inference

We inferred historical changes in effective population size (Ne) for 12 Sisoridae species using the Pairwise Sequentially Markovian Coalescent (PSMC) model(136). For each species, high-quality whole-genome resequencing reads were mapped to their respective reference genomes, and consensus diploid sequences were generated from variant calls using vcfutils.pl (vcf2fq -d 10 -D 80), where sites with read depth lower than 10 or higher than 80 were filtered out. The resulting consensus sequences were converted into PSMC input format using fq2psmcfa. PSMC analyses were then performed with the parameters -N25 -t15 -r5. Default time interval settings were used unless otherwise specified. To scale the results to real time and population size, we applied a mutation rate (μ) of 1.0×10^-8^ per site per generation and a generation time of 4 years, based on estimates from related teleost species.

### Ortholog identification

Orthologous genes were identified across 23 species, including 12 Sisoridae species and 11 outgroup taxa (*Ameiurus mela*s, *Clarias gariepinus*, *Danio rerio*, *Hemibagrus wyckioides*, *Neoarius graeffei*, *Pangasianodon hypophthalmus*, *Silurus asotus*, *Silurus meridionalis*, *Tachysurus fulvidraco*, *Trichomycterus rosablanca*, and *Ictalurus punctatus*), using both clustering- and similarity-based approaches.

First, orthogroups were inferred using OrthoFinder (v2.5.4), yielding a total of 24,350 orthogroups, including 3,121 single-copy orthologs shared across all species(137). In parallel, pairwise orthologs were identified using an iterative reciprocal best hit (RBH) strategy implemented with MMseqs2, using *Ictalurus punctatus* as the reference genome(138). Protein sequences were compared in an all-versus-reference manner with an e-value cutoff of 1e-5, and reciprocal best hits were retained as candidate orthologs. To improve sensitivity, three rounds of RBH searches were performed. In each round, identified RBH pairs were extracted and removed, and the remaining unmatched sequences were subjected to subsequent rounds of similarity searches.

RBH-derived orthologs were further refined using a multi-step filtering strategy. In the first filtering step, gene pairs with sequence identity greater than 50% were retained, resulting in 22,265 orthologous gene pairs, among which 7,096 were present across all 23 species(Supplementary Table 12).

In the second filtering step, additional constraints on alignment coverage and length consistency were applied to ensure orthology reliability. Specifically, three metrics were considered: (i) query coverage (qcov), defined as alignment length divided by query length; (ii) subject length ratio (slen_ratio), defined as subject length divided by query length; and (iii) alignment coverage on the subject (aln_scov), defined as alignment length divided by subject length. Gene pairs satisfying qcov ≥ 0.7, slen_ratio ≥ 0.7, and aln_scov ≥ 0.9 were retained. This step yielded 20,736 orthologs, including 2,730 shared across all 23 species and 3,976 shared across all 12 Sisoridae species. After excluding Bagarius yarrelli due to relatively low annotation quality, 5,905 orthologs were retained across the remaining Sisoridae species.

In the third filtering step, we applied MACSE(v2) to detect and correct potential frameshift mutations(139). For genes showing frameshift signals in Sisoridae species, the corresponding sequences were treated as frameshift-affected and excluded from downstream analyses. In contrast, frameshifts detected only in outgroup species were corrected using MACSE-reconstructed sequences. This filtering step was primarily designed for downstream convergent evolution analyses and resulted in a final dataset of 3,313 high-confidence orthologs.

### Ancestral state reconstruction

To reconstruct the evolutionary history of elevational adaptation in Sisoridae, we mapped species elevation onto a time-calibrated phylogeny using the R package phytools (version 2.0)(140). Elevation data were obtained from published distribution records (Supplementary Table 13). Continuous trait evolution was reconstructed under a maximum likelihood framework assuming a Brownian motion model using the contMap function. This approach estimates ancestral states at internal nodes and interpolates trait values along branches. The resulting trait gradients were visualized using a color ramp from low to high elevation.

### Genome-wide evolutionary rate estimation

To estimate genome-wide evolutionary rates in Sisoridae, we calculated substitution rates (substitutions per site per million years) based on whole-genome alignment (WGA) data. Specifically, substitution rates were defined as the ratio of branch length (substitutions per site) to divergence time. From the WGA dataset, we extracted ∼44.8 Mb of aligned sequences that were shared across all Sisoridae species and contained no gaps. These alignments were partitioned into 1,000 approximately equal-sized segments, and a phylogenetic tree was independently reconstructed for each segment using IQ-TREE2 with model selection (-m MFP). For each inferred tree, root-to-tip branch lengths were calculated for all species after rerooting using a predefined outgroup. To ensure consistency across loci, trees with topologies inconsistent with the reference species tree were excluded from downstream analyses. In total, 692 trees passed the topology filtering and were retained for rate estimation(Supplementary Table 14). Divergence times for each species were obtained from a time-calibrated phylogeny. Evolutionary rates were then calculated for each species and each locus as the ratio of root-to-tip branch length to the corresponding divergence time. The resulting per-locus rate estimates were aggregated to obtain genome-wide distributions of evolutionary rates for each species.

The relationship between evolutionary rate and species elevation was tested using phylogenetic generalized least squares (PGLS) regression implemented in the R package phylolm (https://github.com/lamho86/phylolm). Species present in both the trait dataset and the phylogeny were retained for analysis. Multiple evolutionary models (BM, OU, lambda, kappa, delta, and trend) were fitted, and model fit was evaluated using Akaike Information Criterion (AIC) with corresponding AIC weights. The correlation between evolutionary rate and elevation was assessed based on the fitted PGLS models (Supplementary Table 15). A standard linear regression was also performed as a non-phylogenetic comparison.

Evolutionary rate variation through time in Sisoridae was assessed using internal nodes of a phylogenetic tree constructed from the 44.8 Mb no-gap WGA dataset. For each internal node, the cumulative branch length from the root to the node was calculated from the maximum likelihood tree, and the corresponding divergence time was obtained from a time-calibrated phylogeny. Evolutionary rates were then estimated as the ratio of node-to-root branch length to divergence time (Supplementary Table 16). The relationship between evolutionary rate and time was modeled using locally weighted regression (LOESS), and 95% confidence intervals were estimated using bootstrap resampling. The resulting trends were visualized to illustrate temporal dynamics of evolutionary rates across the Sisoridae phylogeny. Major geological events, including the uplift of the Himalaya and India-Asia collision system (IAS), the Lhasa-Qiangtang suture zone, and Eastern Tibet, were overlaid for comparison(60).

### Lineage-specific acceleration of protein-coding gene evolution

To investigate lineage-specific evolutionary rate variation in Sisoridae, we analyzed 3,976 orthologous genes obtained after stringent filtering. For each Sisoridae species, branch model analyses were performed using the codeml program in PAML, in which each focal Sisoridae species was independently designated as the foreground branch, while available outgroup species were included as background taxa, depending on gene presence across species(114). Model-derived estimates of dN, dS, and dN/dS (ω) were extracted from codeml output for each gene (Supplementary Table 17). Species-specific distributions of dN, dS, and ω were summarized across all genes and visualized using boxplots.

To further characterize gene-level variation in evolutionary rates, ω values were organized into a gene-by-species matrix and normalized using Z-score transformation across genes. The resulting matrix was used to visualize relative rate shifts among species, and hierarchical clustering of genes performed to identify patterns of co-variation(Supplementary Table 18). Genes showing elevated Z-score values in each species were identified and subjected to functional enrichment analysis using Metascape to infer biological processes potentially associated with lineage-specific acceleration of protein-coding gene evolution (Supplementary Table 19). An initial threshold of Z ≥ 2.0 was applied to define candidate genes. For species in which no significant enrichment was detected, the threshold was sequentially relaxed to Z ≥ 1.8 and, if necessary, Z ≥ 1.5, to obtain sufficient candidate genes for downstream enrichment analysis.

### Lineage-specific positive selection and accelerated evolution analyses

To investigate lineage-specific adaptive evolution in Sisoridae, we first defined multiple phylogenetically informed focal groups based on morphological and ecological characteristics, including a full Sisoridae group (all 12 species) and several subgroups within Sisoridae (Fig. 5A). For each analysis, species within a given focal group were designated as foreground lineages, while all remaining Sisoridae species outside the focal group together with available outgroup species were used as background taxa, depending on gene presence in the filtered RBH ortholog dataset.

Positive selection analyses were performed using the branch-site model implemented in the codeml program of the PAML package (v4.8)(114). For each orthologous gene, a likelihood ratio test (LRT) was used to compare the alternative model allowing positive selection at sites on the foreground branches against the corresponding null model. P-values were derived from Chi-square distributions, and genes with nominal LRT P < 0.05 were considered candidate positively selected genes (PSGs) (Supplementary Table 20).

In parallel, lineage-specific acceleration of protein-coding gene evolution was assessed using the branch model in codeml. A null model assuming a single ω ratio across all branches was compared to an alternative model allowing different ω values between foreground and background branches. Genes showing both a significant LRT (P < 0.05) and elevated ω values in the foreground lineage were considered to exhibit accelerated evolution.

### Convergent evolution analysis

To investigate convergent amino acid substitutions in Sisoridae, we analyzed 3,313 high-confidence orthologous genes after stringent frameshift filtering (see above). Protein alignments and corresponding rooted phylogenetic trees were analyzed using the CSUBST pipeline(141). For each orthologous gene, convergent amino acid substitutions were inferred using the csubst analyze module under the ECMK07+F+R4 codon substitution model implemented in IQ-TREE2. Pairwise branch comparisons were restricted to two-lineage configurations (max_arity=2) with exhaustive searches (exhaustive_until=2), and sister-branch comparisons were excluded. The resulting branch-pair comparisons were further filtered to retain high-confidence convergence signals by requiring a node distance of at least three, exclusion of multi-hit artifacts (is_mf=N), elevated convergent ω statistics (omegaCdif2dif>1), positive convergent nonsynonymous substitution signal (dNCdif2dif>0), greater nonsynonymous than synonymous convergence signal (dSCdif2dif < dNCdif2dif), and at least two substitutions on each branch. To focus on convergence among extant Sisoridae species, only terminal branch pairs were retained. These terminal branch pairs were subsequently mapped onto the species tree, and species pairs separated by at least three internodes were classified as distantly related convergent comparisons, whereas more closely related pairs were regarded as lineage-specific substitutions.

Branch-level amino acid substitutions inferred by CSUBST were subsequently reorganized into site-by-species matrices for each gene, in which rows represented amino acid sites and columns represented terminal species with inferred derived amino acid states. Filtered branch pairs were intersected with these matrices to identify candidate site-level substitutions supported by convergent branch-pair evidence. True convergent substitutions were subsequently defined as cases in which two terminal branches independently acquired the same derived amino acid state at the same site. Supported convergent substitutions were then summarized across all genes at the pairwise branch, site, and gene levels. For each site, the number of independent terminal lineages sharing the same derived amino acid state was recorded, and gene-level convergence statistics were calculated by summarizing the number of convergent sites together with the mean and maximum numbers of convergent lineages per site. Genes containing more than 10 convergent sites and at least one site involving four or more independent lineages were retained as high-confidence convergently evolving candidates (Supplementary Table 21). A total of 199 candidate genes were identified, of which 176 annotated protein-coding genes were retained for downstream functional enrichment analysis using Metascape (Supplementary Table 22). In addition, lineage- or group-specific convergent substitutions were identified by restricting analyses to predefined Sisoridae clades, allowing detection of shared amino acid changes within specific evolutionary lineages.

### Analysis of conserved non-coding elements (CNEs)

We expanded our previous whole-genome alignments (WGAs) of 12 Sisoridae species by incorporating additional outgroup taxa, including Ameiurus melas, Clarias gariepinus, Danio rerio, Silurus meridionalis, Tachysurus fulvidraco, and Ictalurus punctatus, to improve phylogenetic representation. Conserved non-coding elements (CNEs) were identified using a combination of PhastCons (v1.3; parameters: expected length = 45, target coverage = 0.3, rho = 0.3) and GERP (parameter: -d 0.01)(142). A neutral model was estimated using phyloFit in the PhastCons package (-EM -precision HIGH -subst-mod REV) based on a fourfold degenerate (4d) site tree. Results from both methods were merged using BEDTools (v2.25.0; -d 10), and elements overlapping annotated coding regions were removed(143). Only non-coding elements ≥30 bp were retained.

For downstream dCNE analysis, CNE multiple alignments in MAF format were first filtered to retain high-quality orthologous alignments. Each CNE alignment was required to contain exactly the species included in the predefined species list, with no missing or additional species. Alignments in which the same species mapped to multiple scaffolds were discarded to avoid potentially ambiguous or non-orthologous matches. We further required a minimum alignment length of 30 bp and a minimum sequence identity of 0.6, calculated as the proportion of fully matched non-gap alignment columns among all informative columns. This filtering step retained 6,433 CNE alignments.

The filtered CNE alignments were then screened for insertion/deletion signals. For each MAF block, gap patterns were quantified for each species, including the longest consecutive gap and the overall gap fraction. CNE blocks were retained for subsequent screening if at least one species contained a consecutive gap of 5 bp or longer. To further enrich for Sisoridae-associated CNE indel candidates, we retained only CNEs in which no indel signal was detected in the designated background/outgroup species (*Clarias gariepinus, Silurus meridionalis, Ictalurus punctatus, Tachysurus fulvidraco*, and *Danio rerio*), while at least one of the remaining Sisoridae species showed an indel signal. This procedure identified 251 candidate CNEs for subsequent group-specific dCNE analyses.

CNE-gene associations were determined by projecting CNE coordinates onto the Danio rerio reference genome (GRCz11). Genes overlapping CNEs (including exonic and intronic regions) or located within ±100 kb upstream or downstream were retained, and all associated genes were kept when multiple matches occurred. In total, 229 CNEs were successfully linked to genes, whereas 22 could not be assigned, resulting in 933 associated genes for downstream analyses (Supplementary Table 23).

### Predicting transcription factor binding motifs

To assess candidate transcription factor binding motifs in the hoxc13a-associated CNE, we extracted the ungapped *Danio rerio* and *Pareuchiloglanis macrotrema* (PMAC) sequences from the multispecies alignment of CNE13. The input sequences were 281 and 277 bp, respectively. Each sequence was scanned independently on both strands with FIMO (MEME Suite v5.5.6) against a locally stored JASPAR motif collection in MEME format (879 motif matrices), using a site-level P-value output threshold of 1×10^-4^. Motif models were classified as detected in PMAC only, D. rerio only, or both when at least one site was reported in the corresponding sequence. For models with multiple hits in one sequence, the site with the smallest P value was used as the representative hit in Supplementary Table 24; all reported hits are provided in its accompanying “FIMO hits” worksheet. Hit coordinates are one-based positions within the ungapped input sequences. FIMO-reported q values are included for reference but were not used as an additional filter.

### Visualization of the candidate CNE near rgma and chd2

The zebrafish coordinate of the candidate element was converted from danRer11 chr18:24,995,683 – 24,995,724 to danRer10 chr18:25,009,151 – 25,009,192 using UCSC liftOver and the danRer11ToDanRer10 chain file. For locus visualization, we obtained previously processed adult Danio rerio muscle RNA-seq, ATAC-seq, H3K27ac, H3K4me3, and Hi-C tracks from the zebrafish epigenome atlas of Yang et al. (GEO: GSE134055). The published bigWig signal tracks were plotted directly; we did not realign their sequencing reads or call peaks. The downloaded muscle Hi-C .hic file was converted to 5-kb and 10-kb .cool matrices with hic2cool and balanced with cooler. Insulation scores were calculated from the 5-kb matrix using cooltools with 250-kb and 500-kb windows and the Li boundary threshold. The 5-kb contact map, 250-kb insulation track, gene annotations, and published signal tracks were displayed with figeno using the danRer10 reference. The Hi-C view covered chr18:24,890,000 – 25,150,000, and the RNA-seq and epigenomic view covered chr18:24,930,000 – 25,030,000. These analyses were used to visualize the zebrafish locus; enhancer activity and target-gene regulation in Sisoridae were not tested.

## Supporting information

Supplementary Table 6. QuIBL results for all species triplets based on chromosome-wise analyses

Supplementary Table 10. Genes associated with introgression identified by AutoCoalHMM at node7

Supplementary Table 11. Tissue expression profiles of V2-type introgression-associated genes

Supplementary Table 12. Orthologous gene assignments with RBH method

Supplementary Table 17. Lineage-specific gene-wise dN, dS and dNdS estimates inferred by PAML branch models for 12 Sisoridae species

Supplementary Table 19. Functional enrichment results for lineage-specific genes exhibiting elevated evolutionary rate Z-scores

Supplementary Table 20. Complete positive selection results for different adaptive trait groups in Sisoridae

Supplementary Table 21. Genome-wide summary of convergent amino acid substitutions identified by CSUBST

Supplementary Table 22. Functional enrichment analysis of high-confidence convergently evolving genes

Supplementary Table 24. Predicted TF motifs in CNE13

## Acknowledgements

This work was supported by the National Natural Science Foundation of China (32422010, 32500392) and the National Natural Science Foundation of China (NSFC) Joint Fund Priority Support Program (U23A20249). Thanks for all members of the Research Group on Fish Phylogenetics and Adaptive Evolution for collecting the samples.

## Author contributions

S. H. and Y.C. lead this study. L.Y. conceived and supervised the study. Y.Q., M.M., Z.D. analyzed the data. T.M., W.C., Y.L., L.Z., C.W., and X.G. sampled the data. Y.Q. and L.Y. wrote the paper with input from all authors.

## Competing interests

The authors declare no competing interests.

## Notes

### Competing Interest Statement

The authors have declared no competing interest.

