## Supplementary Figures for "The uplift of the Qinghai-Tibetan Plateau reshaped genome architecture and diversification of Asian torrent catfishes"

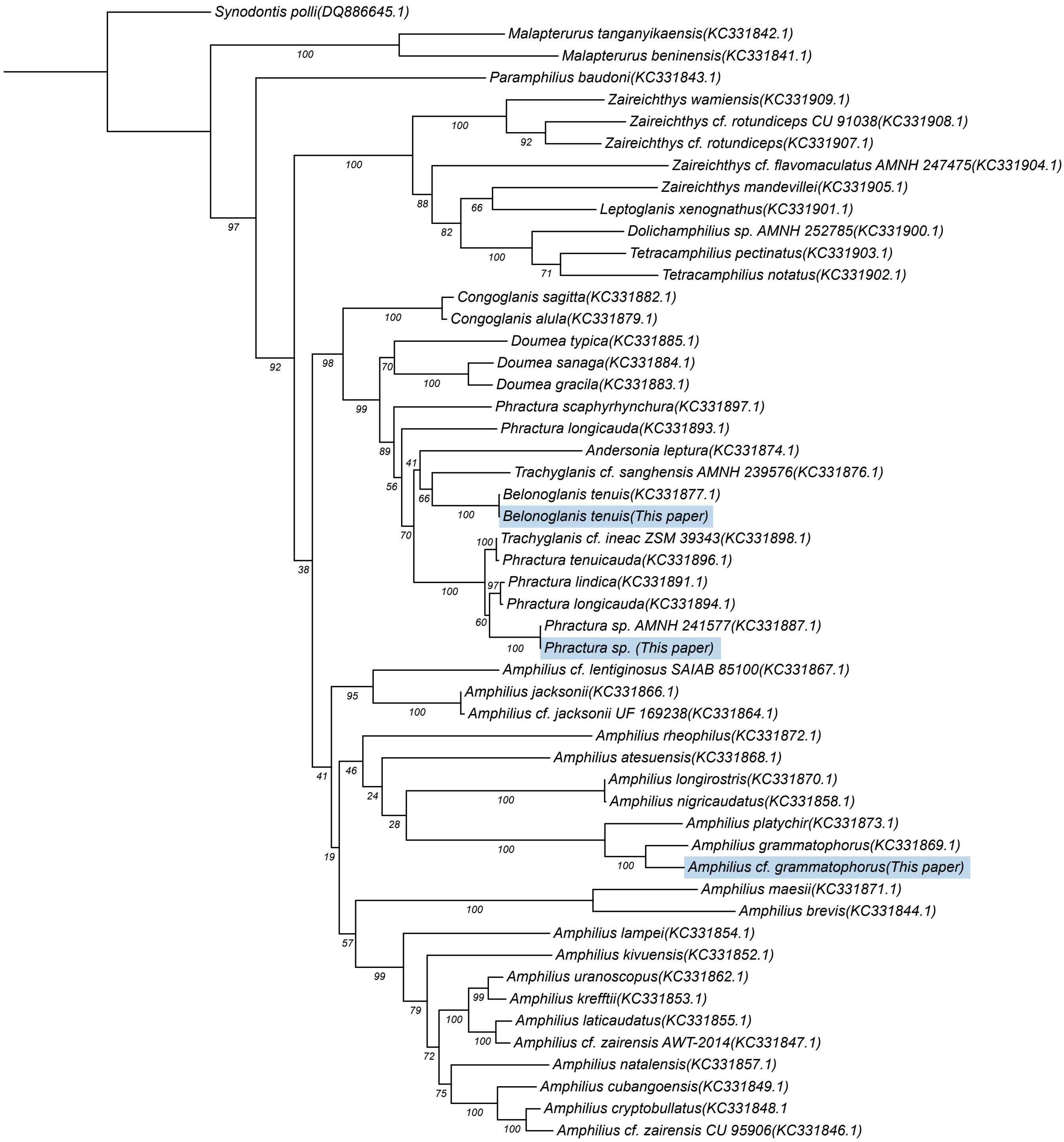


**Supplementary Fig 1. Species identification of Amphiliidae specimens.**
Maximum-likelihood phylogeny of African siluriform fishes reconstructed from cytochrome b (Cytb) gene sequences. The three Amphiliidae specimens sampled in this study were assigned to Belonoglanis tenuis, Phractura sp., and Amphilius cf. grammatophorus, respectively. The number following each species represents the NCBI ACCESSION and VERSION numbers for the corresponding gene.

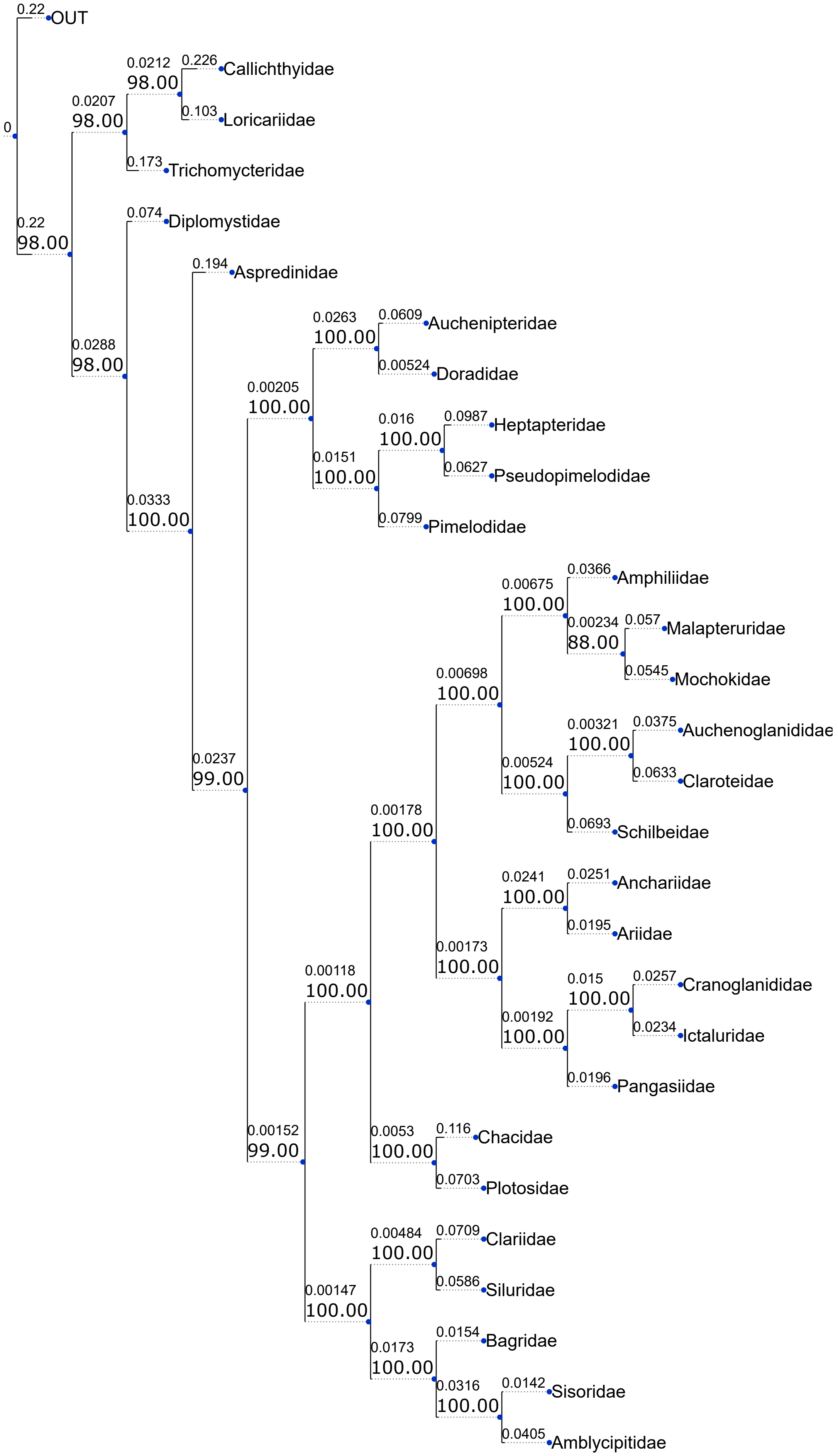


**Supplementary Fig 2. Maximum-likelihood phylogeny inferred from the full codon dataset.**
Maximum-likelihood (ML) phylogeny of Siluriformes reconstructed from the concatenated alignment of all 1,896 BUSCO genes using full codon sequences. For visualization purposes, terminals were collapsed to the family level. Corresponding species-level phylogenies are provided in Supplementary Data 1. Numbers at nodes indicate ultrafast bootstrap support values.

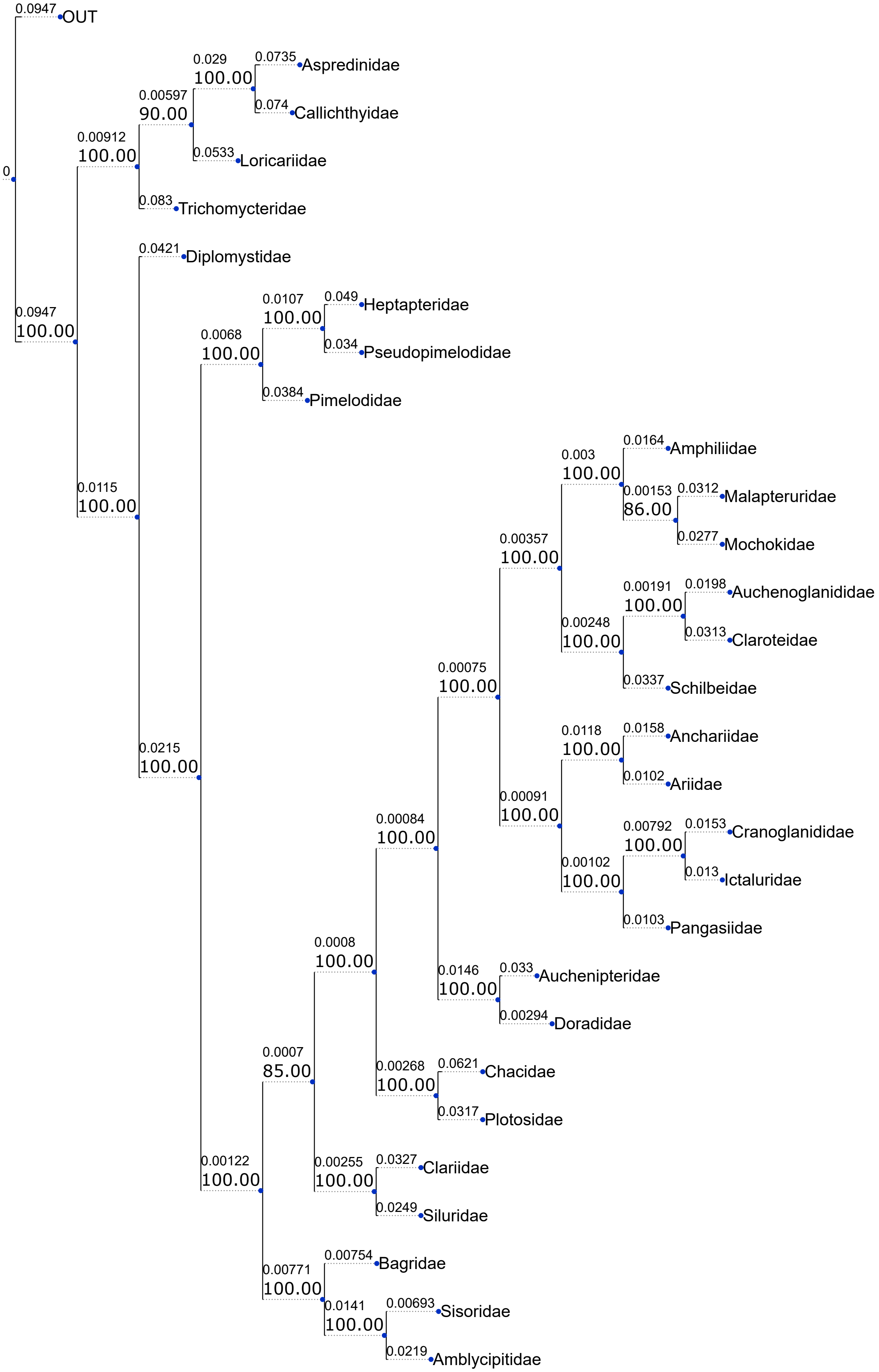


**Supplementary Fig 3. Maximum-likelihood phylogeny inferred from the first and second codon positions (P1+P2).**
ML phylogeny reconstructed from the concatenated alignment of the first and second codon positions (P1+P2) of 1,896 BUSCO genes. For visualization purposes, terminals were collapsed to the family level. Corresponding species-level phylogenies are provided in Supplementary Data 1. Numbers at nodes indicate ultrafast bootstrap support values.

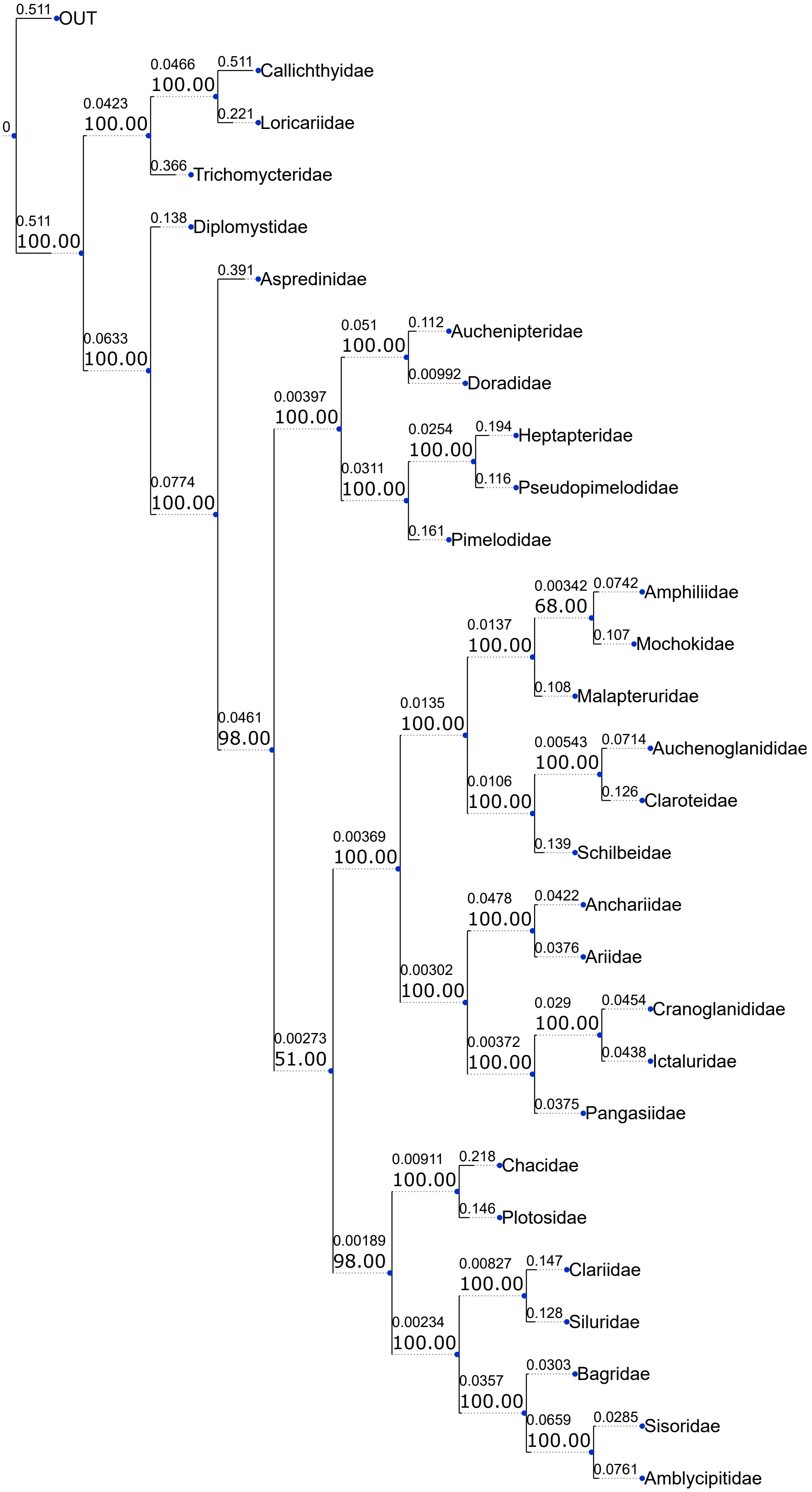


**Supplementary Fig 4. Maximum-likelihood phylogeny inferred from the third codon positions (P3).**
ML phylogeny reconstructed from the concatenated alignment of the third codon positions (P3) of 1,896 BUSCO genes. For visualization purposes, terminals were collapsed to the family level. Corresponding species-level phylogenies are provided in Supplementary Data 1. Numbers at nodes indicate ultrafast bootstrap support values.

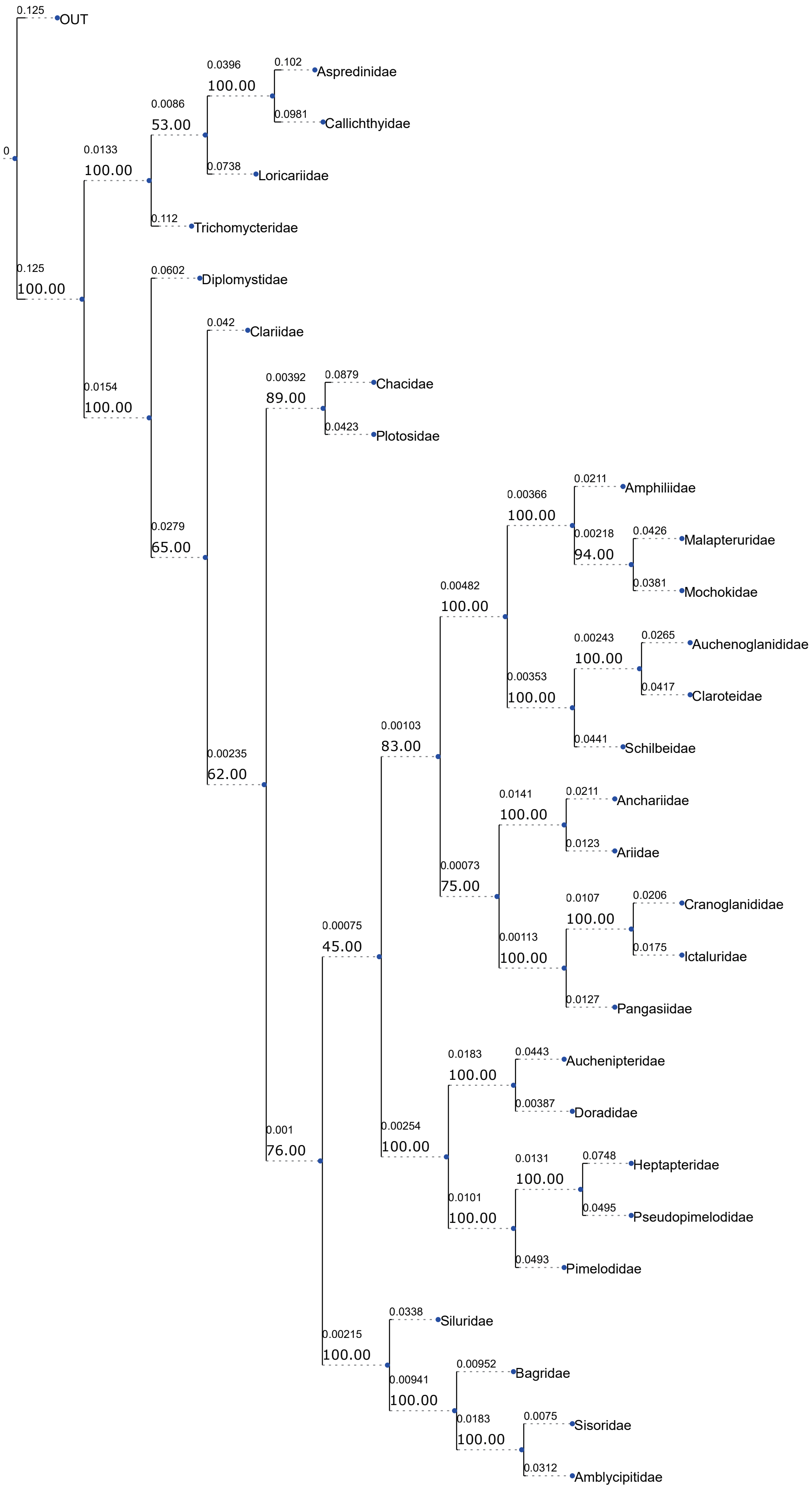


**Supplementary Fig 5. Maximum-likelihood phylogeny inferred from amino acid sequences.**
ML phylogeny reconstructed from the concatenated amino acid alignment of 1,896 BUSCO genes. For visualization purposes, terminals were collapsed to the family level. Corresponding species-level phylogenies are provided in Supplementary Data 1. Numbers at nodes indicate ultrafast bootstrap support values.

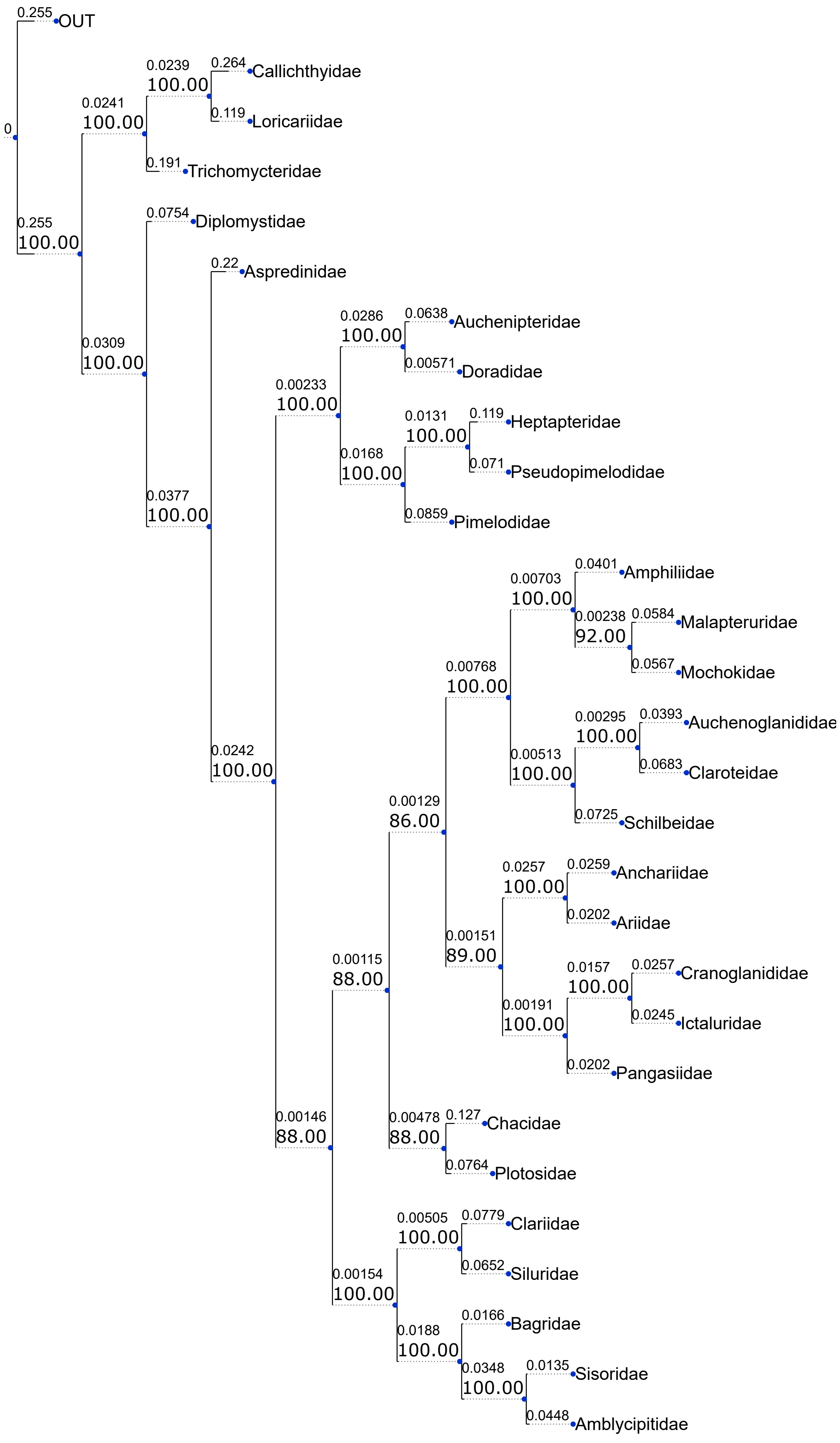


**Supplementary Fig 6. Maximum-likelihood phylogeny inferred from the fast-evolving BUSCO genes.**
ML phylogeny reconstructed from the concatenated alignment of the fastest-evolving 50% of BUSCO genes, ranked according to evolutionary rates obtained from the OrthoDB database. For visualization purposes, terminals were collapsed to the family level. Corresponding species-level phylogenies are provided in Supplementary Data 1. Numbers at nodes indicate ultrafast bootstrap support values.

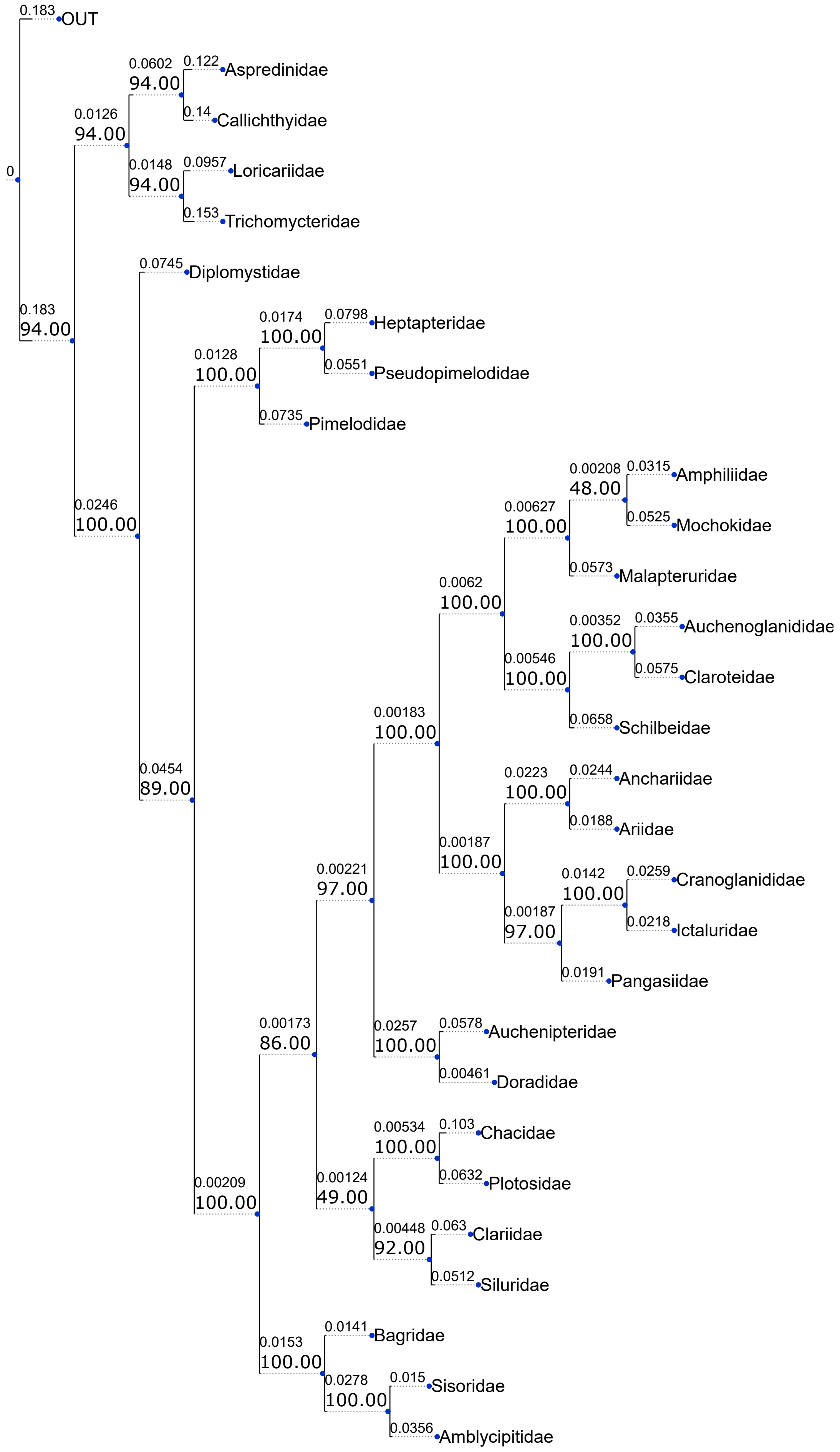


**Supplementary Fig 7. Maximum-likelihood phylogeny inferred from the conserved BUSCO genes.**
ML phylogeny reconstructed from the concatenated alignment of the most conserved 50% of BUSCO genes, ranked according to evolutionary rates obtained from the OrthoDB database. For visualization purposes, terminals were collapsed to the family level. Corresponding species-level phylogenies are provided in Supplementary Data 1. Numbers at nodes indicate ultrafast bootstrap support values.

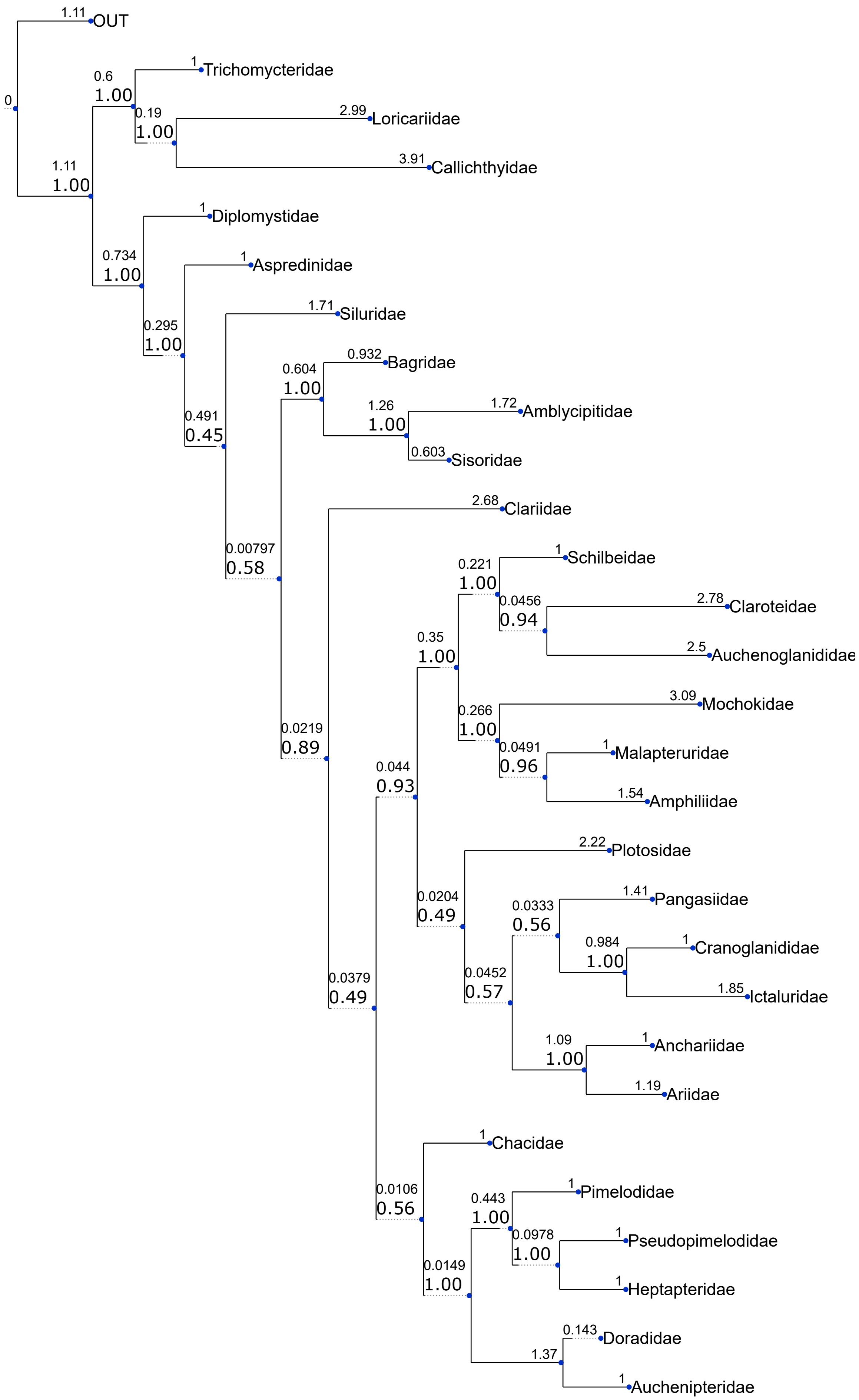


**Supplementary Fig 8. Coalescent-based species tree inferred under the multi-species coalescent model.**
Species tree inferred from 1,896 individual BUSCO gene trees using the coalescent-based method implemented in ASTRAL under the multi-species coalescent (MSC) framework. For visualization purposes, terminals were collapsed to the family level. The corresponding species-level phylogeny is provided in Supplementary Data 1. Numbers at nodes indicate local posterior probabilities (LPPs).

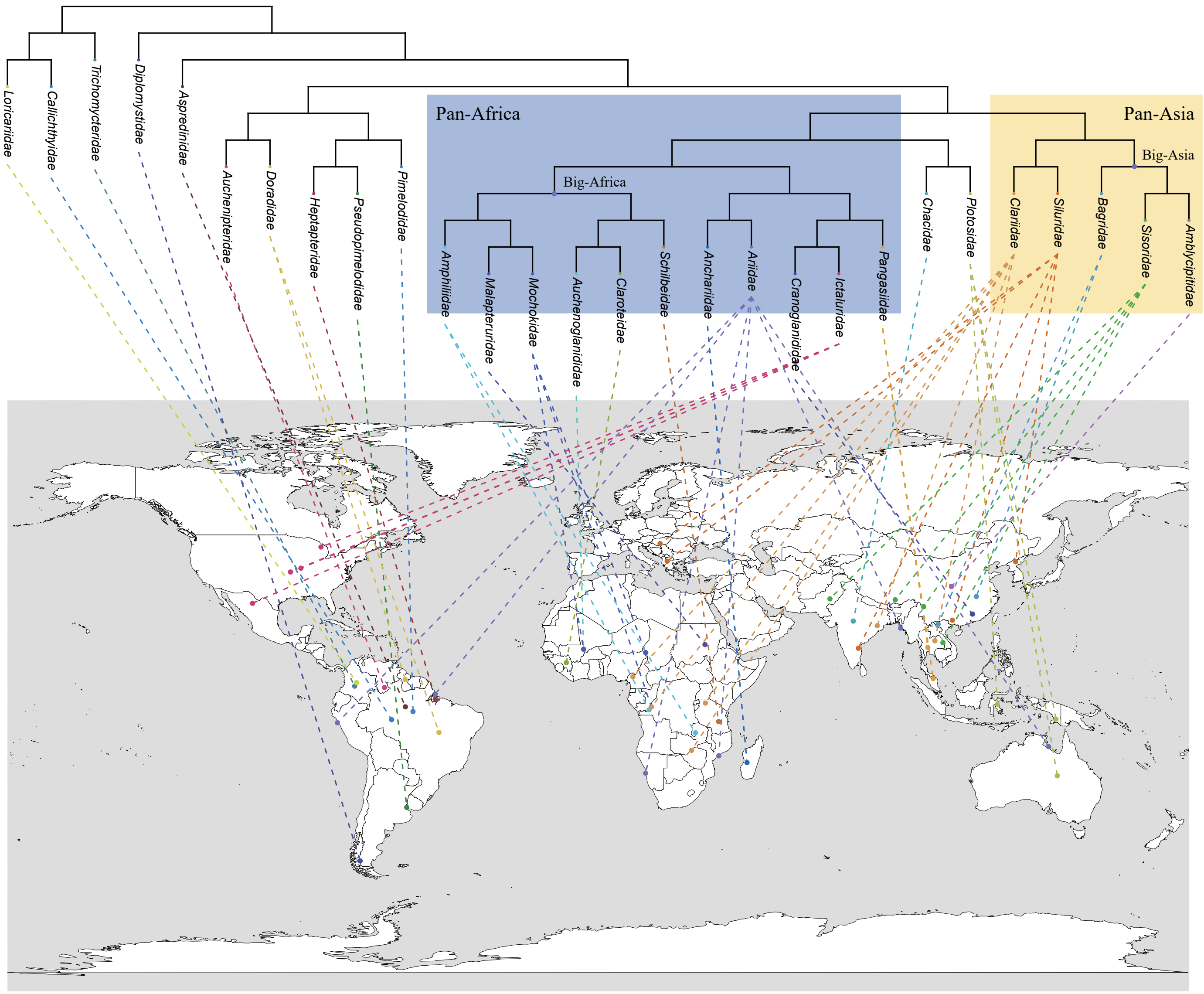


**Supplementary Fig 9. Geographic distribution of representative species across Siluriformes families.**
Map showing the geographic distribution of representative species from each Siluriform family included in this study. Connecting-line colors correspond to the branch colors of the respective families in Figure 1. The major phylogenetic assemblages (Pan-Asia, Big Asia, Pan-Africa, and Big Africa) are highlighted.

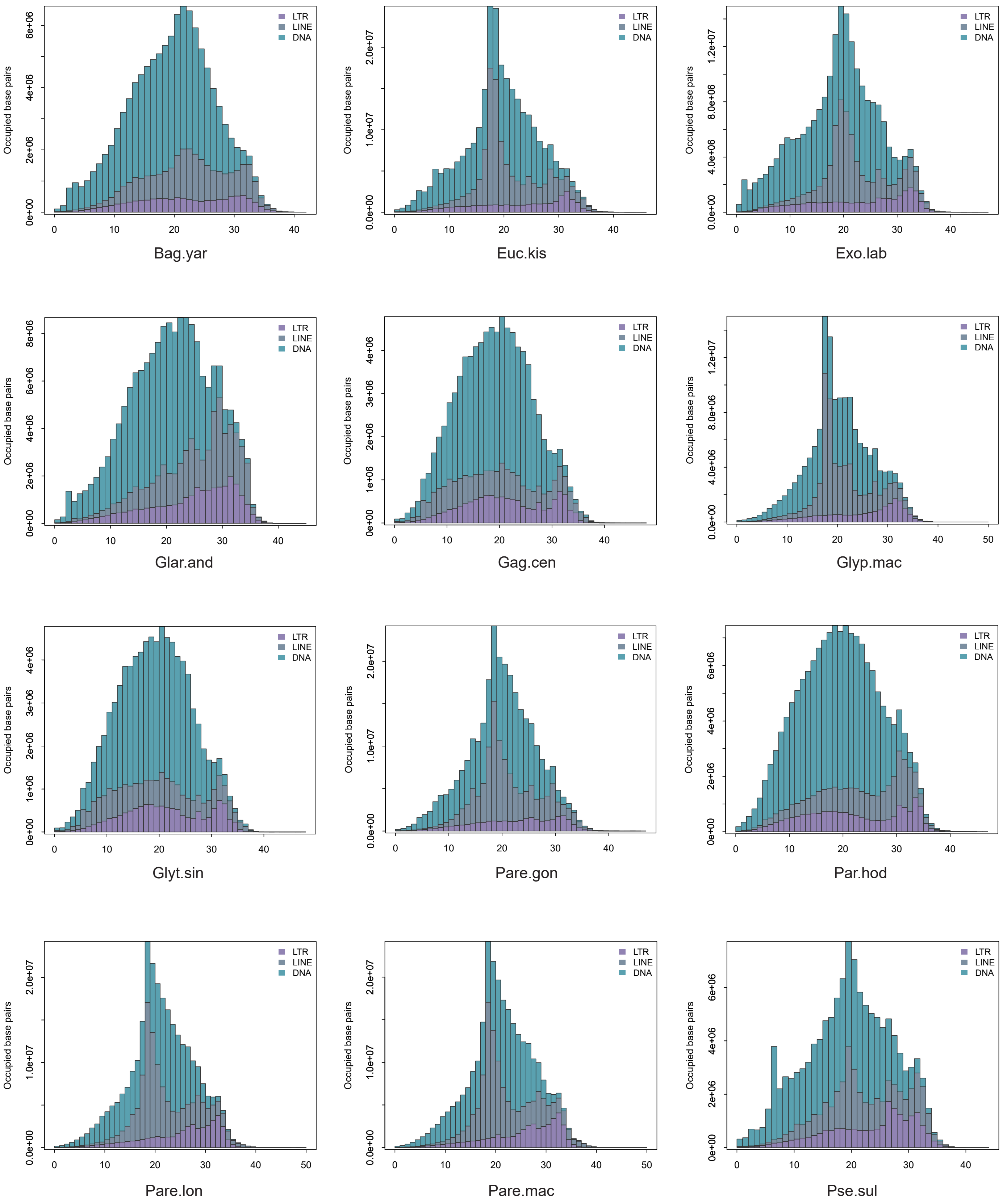


**Supplementary Fig 10. Transposable element landscapes of Sisoridae species.**
Transposable element (TE) landscape plots of Sisoridae species generated using Kimura distance-based copy divergence analyses from RepeatMasker annotations. The x-axis represents Kimura substitution levels, and the y-axis indicates the genomic proportion occupied by TE sequences. Colors denote different TE classes.

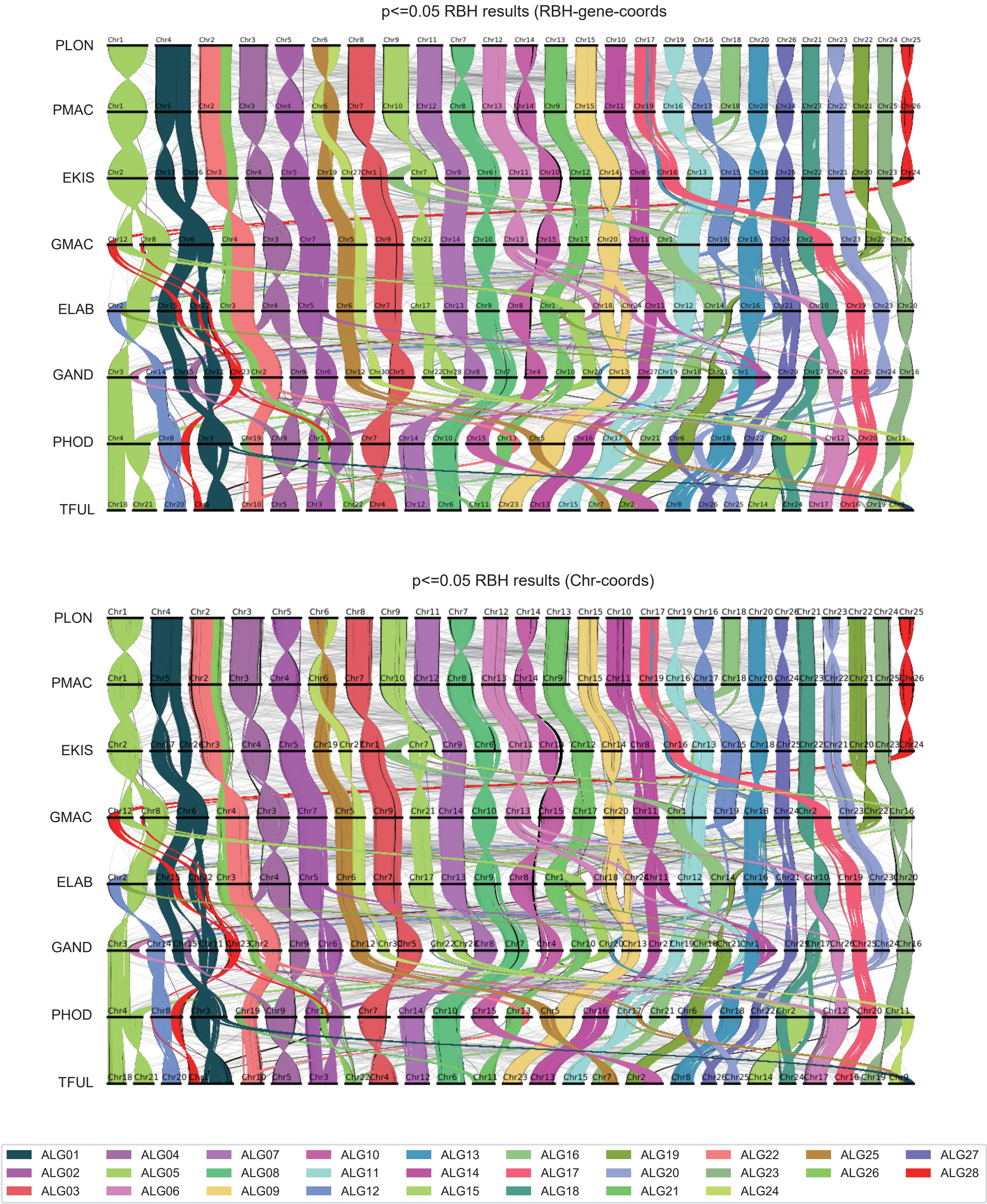


**Supplementary Fig 11. Whole-genome synteny among chromosome-level Sisoridae genomes based on ancestral linkage groups.** Whole-genome synteny comparisons among seven chromosome-level genomes from Sisoridae. Chromosomes are colored according to their assigned ancestral linkage groups (ALGs), with the same color indicating the same ALG. The upper panel shows syntenic relationships identified using reciprocal best-hit (RBH) genes mapped to gene coordinates (RBH-gene-coords; P ≤ 0.05), whereas the lower panel shows syntenic relationships based on chromosome coordinates (RBH-Chr-coords; P ≤ 0.05).

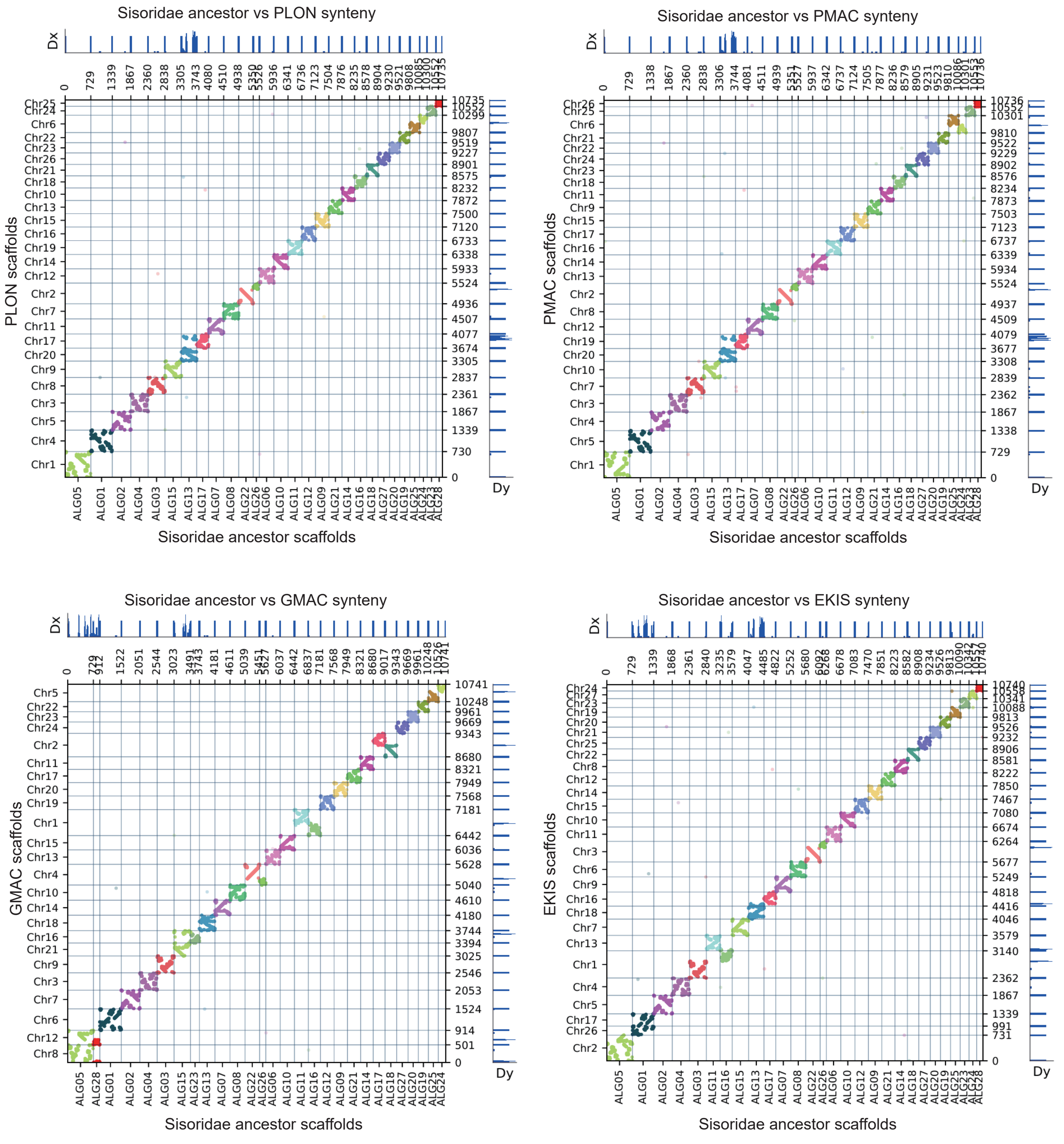


**Supplementary Fig 12. Conserved synteny between the reconstructed Sisoridae ancestral genome and extant Sisoridae genomes.** Dot-plot comparisons between the reconstructed Sisoridae ancestral chromosomes and eight chromosome-level Sisoridae genomes. The x-axis represents chromosomes of each extant genome and the y-axis represents reconstructed ancestral chromosomes. Each dot corresponds to a reciprocal best-hit (RBH) orthologous gene pair. Dot opacity reflects the significance of syntenic associations based on Fisher's exact test, with opaque dots indicating significant associations ($P \leq 0.05$). Dx and Dy profiles summarize changes in syntenic continuity along the x- and y-axes, respectively. Panels (a–h) correspond to the eight chromosome-level Sisoridae genomes analyzed in this study.

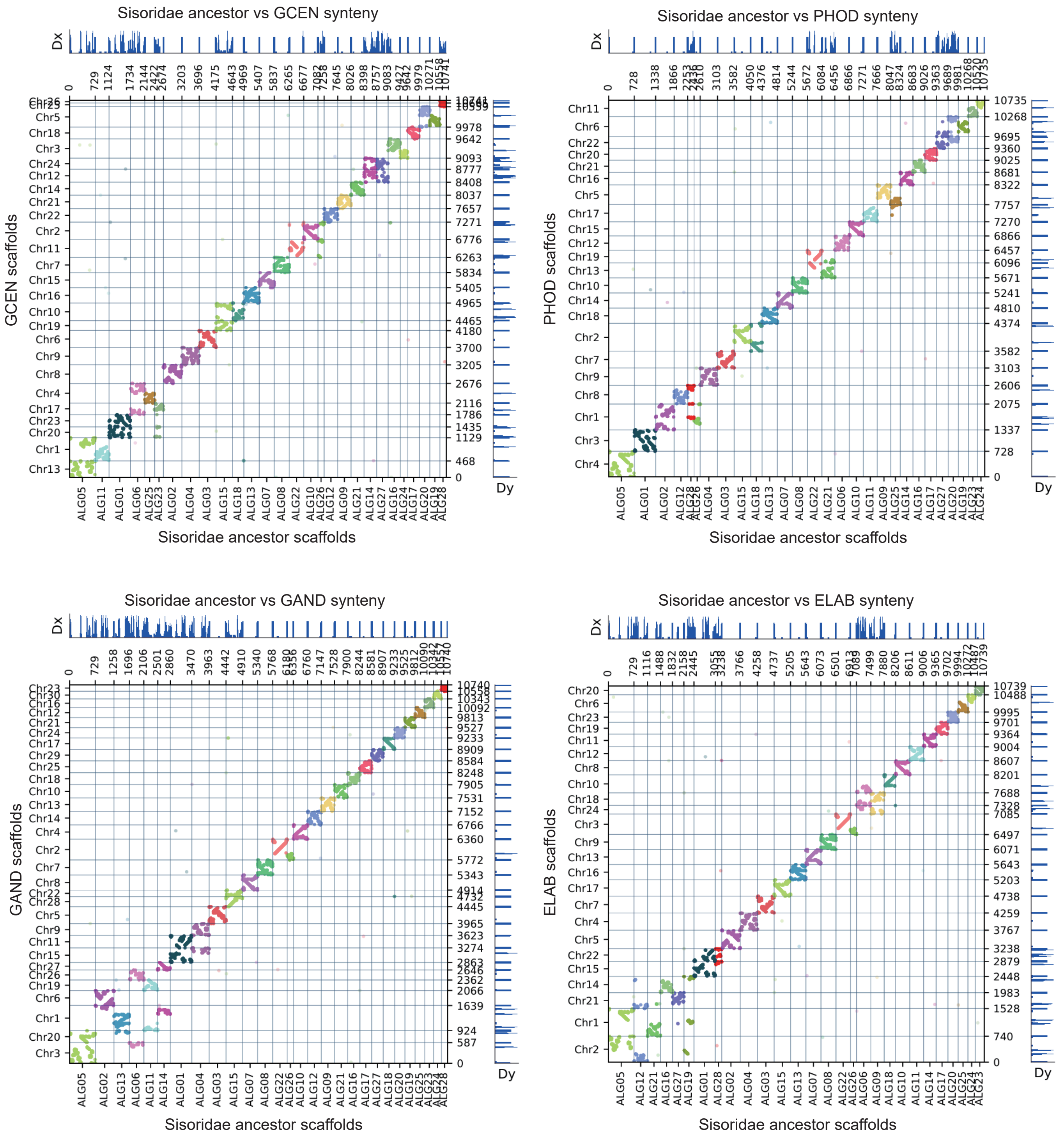
Sisoridae ancestor vs GCEN synteny
GCEN scaffolds
Sisoridae ancestor scaffolds
Dx
Dy
Sisoridae ancestor vs PHOD synteny
PHOD scaffolds
Sisoridae ancestor scaffolds
Dx
Dy
Sisoridae ancestor vs GAND synteny
GAND scaffolds
Sisoridae ancestor scaffolds
Dx
Dy
Sisoridae ancestor vs ELAB synteny
ELAB scaffolds
Sisoridae ancestor scaffolds
Dx
Dy

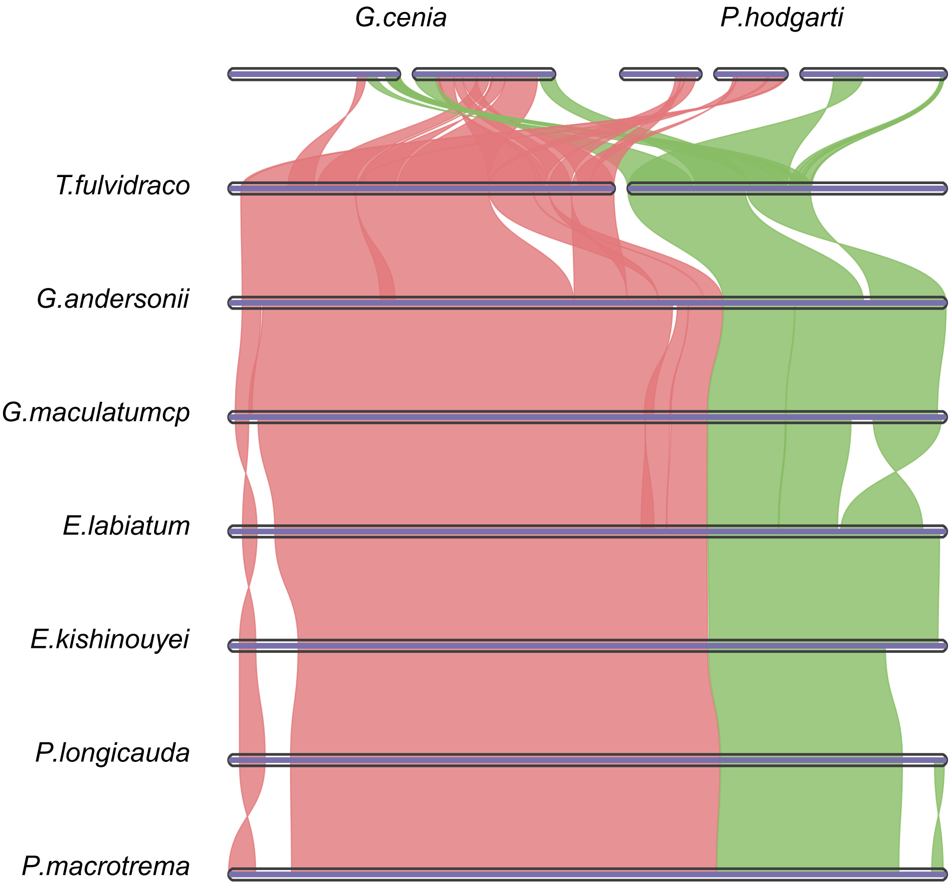


**Supplementary Fig 13. Conserved fusion of ancestral linkage groups ALG22 and ALG16 across Sisoridae.**
Syntenic relationships illustrating the evolutionary history of the fusion between ancestral linkage groups (ALGs) 22 and 16 in eight chromosome-level Sisoridae genomes relative to the non-sisorid catfish Tachysurus fulvidraco. Colored ribbons represent orthologous chromosomal segments corresponding to ALG22 (red) and ALG16 (green). In T. fulvidraco, the two ALGs are retained as separate chromosomes, whereas they are fused into a single chromosome in most Sisoridae species, indicating that this chromosomal fusion represents a conserved structural feature during Sisoridae evolution. The basal lineages Gagata cenia and Pareuchiloglanis hodgarti retain the unfused ancestral configuration.

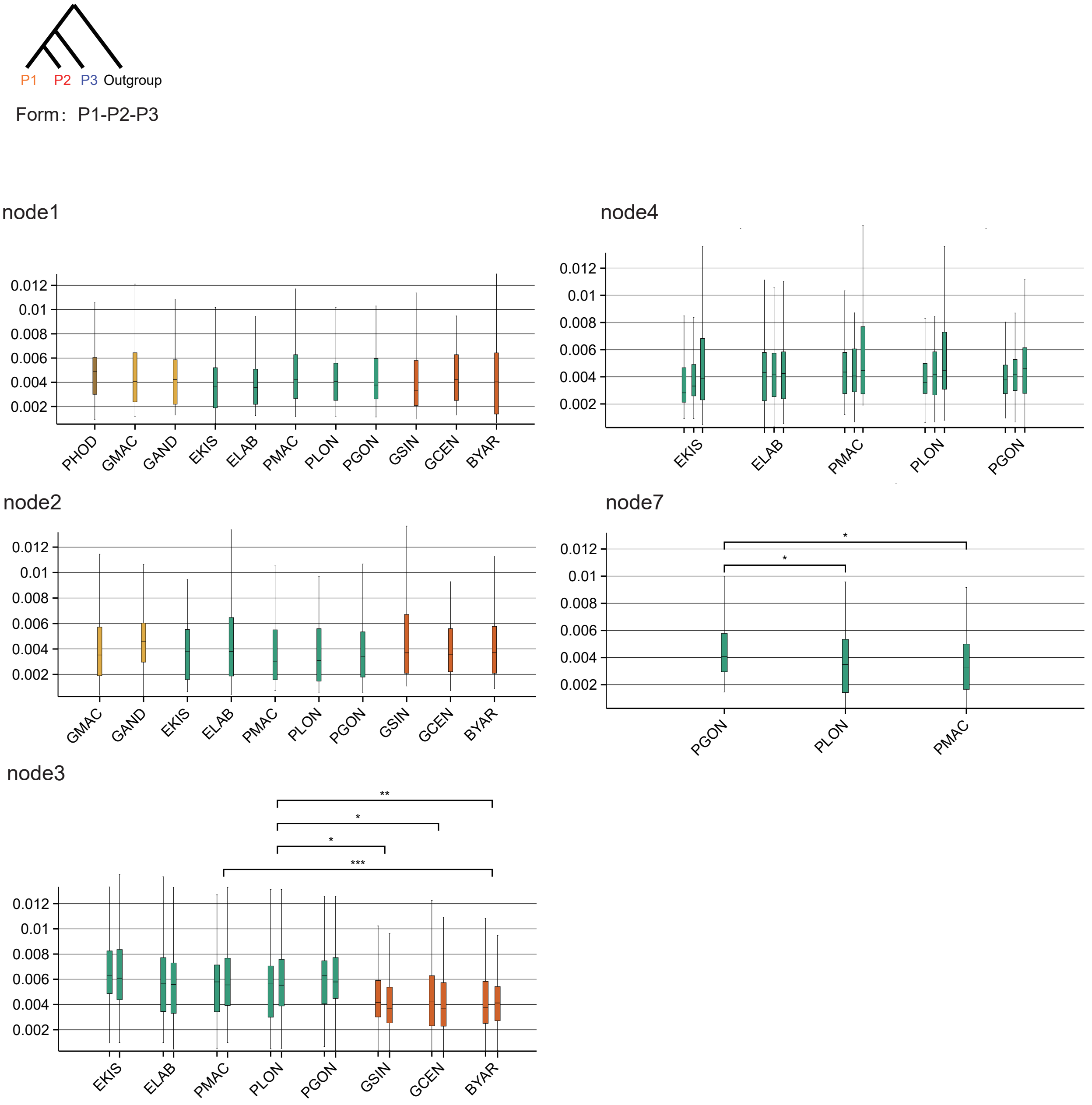


**Supplementary Fig 14. Estimated introgression proportions (totalIntroProp) across introgression-dominated nodes inferred by QuIBL.**

Top, schematic illustration of the QuIBL triplet configuration used in this study. Each QuIBL analysis was performed on a three-taxon configuration (P1–P2–P3), in which P1 and P2 represent sister lineages in the species tree and P3 represents the outgroup of the focal node. Bottom, distributions of the estimated introgression proportion (totalIntroProp) for the five introgression-dominated nodes (nodes 1–4 and node 7). For each node, boxplots summarize totalIntroProp values inferred from all QuIBL triplets sharing the same potential introgressing lineage pair. For nodes containing multiple descendant taxa within a lineage, separate boxplots represent different species combinations satisfying the corresponding triplet configuration. Boxes indicate the interquartile range (IQR), horizontal lines indicate the median, whiskers extend to 1.5 × IQR, and significance between groups was assessed using pairwise statistical comparisons ($P < 0.05$, $P < 0.01$, $P < 0.001$).

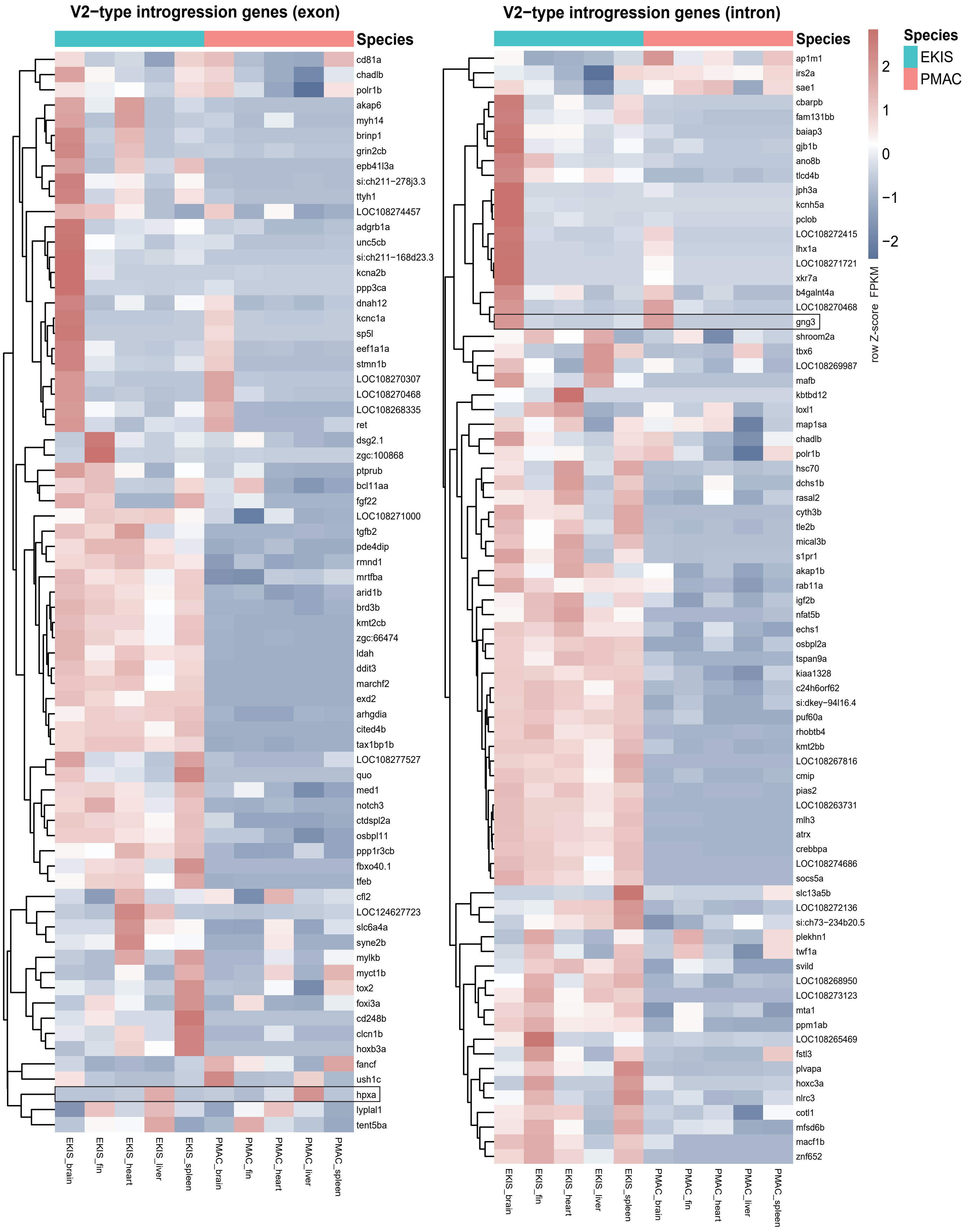


**Supplementary Fig 15. Expression patterns of exon- and intron-defined V2-type introgression-associated genes.**
Heatmaps showing tissue-specific expression patterns of V2-type introgression-associated genes identified from (left) exonic sites and (right) intronic sites in the AutoCoalHMM analysis. V2-type genes correspond to loci supporting the discordant EKIS–PMAC-like genealogy. Gene expression levels (FPKM) were log-transformed and row-scaled (Z-score) for visualization. Columns represent corresponding tissues from E. kishinouyei (EKIS) and P. macrotrema (PMAC), and rows represent individual genes.

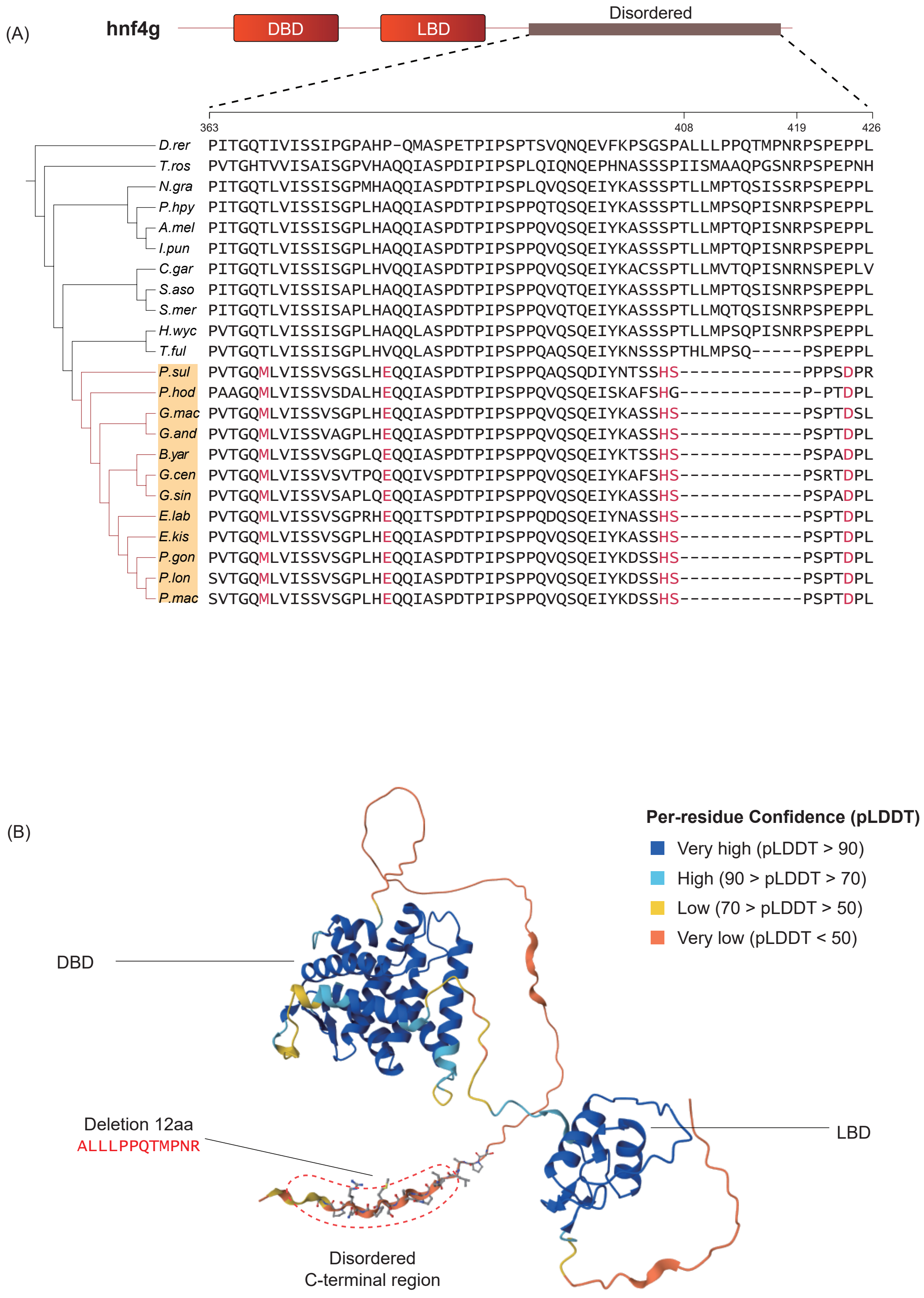


**Supplementary Fig 16. Multiple sequence alignment and AlphaFold2-predicted structure of HNF4g.**
(A) Multiple sequence alignment of representative teleost HNF4g proteins showing a Sisoridae-specific 12-amino-acid deletion within the C-terminal region. Conserved residues are highlighted, and the lineage-specific deletion is indicated by dashed gaps. The domain architecture of HNF4g, including the DNA-binding domain (DBD), ligand-binding domain (LBD), and intrinsically disordered C-terminal region, is shown above the alignment.
(B) AlphaFold2-predicted structure of zebrafish HNF4g (UniProt: A0A8M6YY37) colored according to per-residue prediction confidence (pLDDT). The segment corresponding to the Sisoridae-specific deletion is highlighted in magenta. The deleted region is located within the predicted intrinsically disordered C-terminal region outside the conserved DNA-binding and ligand-binding domains.
